# Comprehensive characterization of tumor microenvironment in colorectal cancer via histopathology-molecular analysis

**DOI:** 10.1101/2023.02.16.528849

**Authors:** Xiangkun Wu, Hong Yan, Mingxing Qiu, Xiaoping Qu, Jing Wang, Shaowan Xu, Yiran Zheng, Minghui Ge, Linlin Yan, Li Liang

**Author notes:** Correspondence: Li Liang. These authors have contributed equally to this work and share first authorship (Xiangkun Wu, Hong Yan, Mingxing Qiu, Xiaoping Qu). Email: Xiangkun Wu; Hong Yan; Mingxing Qiu; Xiaoping Qu; Jing Wang; Shaowan Xu; Yiran Zheng; Minghui Ge; Linlin Yan.

## Abstract

**Purpose:** To explain how the tumor microenvironment (TME) contributes to biological and clinical heterogeneity of colorectal cancer (CRC).

**Methods:** Using multi-omics analysis, single cell transcriptomic sequencing analysis and artificial intelligence-enabled spatial analysis of whole-slide images, we performed a comprehensive characterization of TME in colorectal cancer (CCCRC).

**Results:** CRC samples were classified into four CCCRC subtypes with distinct TME features, namely, C1 as the proliferative subtype with low immunogenicity; C2 as the immunosuppressed subtype with the terminally exhausted immune characteristics; C3 as the immune-excluded subtype with the distinct upregulation of stromal components and a lack of T cell infiltration in tumor core; and C4 as the immunomodulatory subtype with the remarkable upregulation of anti-tumor immune components. The four CCCRC subtypes had distinct histopathological and molecular characteristics, therapeutic efficacy, and prognosis. The C1 subtype was more sensitive to chemotherapy, the C2 and C3 subtypes were more sensitive to WNT pathway inhibitor SB216763 and Hedgehog pathway inhibitor vismodegib, and the C4 subtype was suitable for ICB treatment. Finally, we established a single-sample gene classifier for identifying the CCCRC subtypes.

**Conclusions:** Our integrative analyses ultimately established a holistic framework to thoroughly dissect the TME of CRC, and the CCCRC classification system with high biological interpretability might facilitate biomarker discoveries and clinical treatment decisions in the future.

## Introduction

Colorectal cancer (CRC) is the third most deadly malignancy worldwide (1), and the incidence of early-onset CRC is steadily increasing (2). CRC at early and localized stages is primarily a preventable and curable disease, but up to 50% of patients with locally advanced disease eventually develop mCRC (3, 4). Therefore, the clinical systematic management of CRC patients is still an unmet medical challenge (4).

With the development of high-throughput technologies and bioinformatics strategies, multi-omics data are used to identify and characterize the molecular subtypes of CRC, such as genomics (5), transcriptomics (6–11) and proteomics (12). The consensus molecular subtype (CMS) integrates six independent classification systems based on transcriptomics; however, it is still not explicitly used to guide clinical treatment (13). TCGA and CPTAC colorectal studies have dissected the molecular heterogeneity of CRC by integrating multi-omics data (14, 15). Nevertheless, multi-omics data are complex and highly dimensional, and extracting valuable information from these data to guide clinical treatment is still a tremendous challenge (16). By reviewing the biological characteristics of the tumor, useful information can be screened for identifying molecular subtypes.

The tumor cells can interact with cellular or non-cellular components, triggering dramatic molecular, cellular and physical changes in the tumor microenvironment (TME) to build a self-sustainable tumor ecosystem (17, 18). Simultaneously, TME profoundly affects tumor biology, responses to therapy, and clinical outcomes, which is a dynamic network mainly comprised of immune components and stromal components (19–21). Furthermore, TME can adversely affect the metabolic activities of tumor, immune and stromal cells, and form diverse metabolic phenotypes (22, 23). Identifying the components of the TME and their functions, as well as the crosstalk between tumor cells and TME contributes to our understanding of the clinical heterogeneity of CRC, thereby bringing about new advances in precision medicine. Previous studies have used immune or stromal components of the TME, or a combination of both, to study the TME (24, 25), but they are insufficient to completely reconstruct the heterogeneity of the TME.

In this study, we considered the tumor cells and its TME as a whole and performed a comprehensive characterization of TME in colorectal cancer (CCCRC), including the functional states of the tumor cells, immune and stromal signatures, and metabolic reprogramming features. We successfully identified the four CCCRC subtypes based on 61 TME-related signatures. Integrated analyses determined that the CCCRC subtypes had distinct histopathological and molecular characteristics, therapeutic efficacy, and prognosis.

## Materials and Methods

A total of 2195 samples were obtained from ten publicly available datasets (**Supplementary Table1**). The eight microarray datasets based on the same platform GPL570 (GSE13067, GSE13294, GSE14333, GSE17536, GSE33113, GSE37892, GSE38832 and GSE39582 datasets) were combined as CRC-AFFY cohort to determine molecular classification. The two RNA sequencing datasets (TCGA and CPTAC datasets) were combined as CRC-RNAseq cohort to validate molecular classification.

After reviewing previously published studies, the Molecular Signatures Database (MSigDB; http://www.gsea-msigdb.org/gsea/msigdb/index.jsp), and the Reactome pathway portal (https://reactome.org/PathwayBrowser/), we obtained 61 signatures related to tumor, immune, stromal, and metabolic reprogramming features (**Supplementary Table2**). Gene set variation analysis (GSVA) was performed to calculate the 61 TME-related signature scores based on gene expression profiles (GEP). We devised a novel molecular classification, called CCCRC, using consensus clustering method (26) based on the 61 TME-related signature scores in the CRC-AFFY cohort. To verify the repeatability and robustness of CCCRC, we used the “pamr.predict” function of the R package “pamr” (27) to classify the CRC samples based on the TME-related signature scores in the CRC-RNAseq cohort.

More details of histopathological examination, multi-omics analysis, scRNA-seq analysis, development of treatment strategies, and statistical analysis are provided in the supplementary material and methods.

## Results

### Establishment of the TME panel

The molecular and clinical features of a tumor are characterized by the functional states of tumor cells, as well as other TME-related signatures, including immune and stromal components, and metabolic reprogramming signatures. In brief, 14 signatures (including angiogenesis, apoptosis, cell cycle, differentiation, DNA damage, DNA repair, EMT, hypoxia, inflammation, invasion, metastasis, proliferation, quiescence, and stemness) were used to describe the functional states of tumor cells. As for the immune signatures, we focused on eight categories of immune cells (T cells, natural killer cells, dendritic cells, macrophages, myeloid-derived suppressor cells, B cells, mast cells, neutrophils) and their subpopulations, as well as the other immune-related signatures. In addition to the signatures of endothelial cells, mesenchymal cells, and the extracellular matrix, we included signatures of cancer stem cells and interactions of cells with the extracellular matrix to characterize the stromal compartments. A total of 7 major metabolic pathways (Amino acid, Nucleotide, Vitamin cofactor, Carbohydrate, TCA cycle, Energy, and Lipid metabolism) were used to reveal the metabolic reprogramming of the TME. According to the above biological framework, a total of 61 TME-related signatures were collected to form the TME panel (**Supplementary Table2**), which ultimately established a holistic approach to thoroughly dissect the TME of CRC.

We used GSVA to calculate the TME-related signature scores for each sample in each cohort. Principal coordinate analysis (PCOA) revealed that the CRC samples could be distinguished from normal samples by the TME-related signatures in the GSE39582 and TCGA cohorts (**Fig. S1A**). We further focused on the signatures of the functional states of tumor cells and cancer stem cells, which could classify CRC and normal samples (**Fig. S1B**). The *P*-values for intercomparisons of the euclidean distances between normal and CRC samples were all <0.05 using PERMANOVA test. Most immune signatures had higher GSVA scores in the normal samples compared with the CRC samples (**Fig. S1C, D**), while stromal signatures and the signatures of the functional states of tumor cells had higher GSVA scores in CRC tissues (**Fig. S1C, D**). As expected, amino acid, carbohydrate, and nucleotide metabolic processes were more prominent in CRC samples, which was consistent with the hallmark of infinite proliferation of tumor cells (**Fig. S1C, D**).

Pearson’s correlation analysis of the TME-related signatures revealed three major patterns bound by positive correlations in the CRC-AFFY cohort (**Fig. S1E**). One pattern defining the proliferation of tumor cells consisted of cell cycle and metabolic reprograming signatures. The second was mainly comprised of immune components, such as T cells, NK cells, MDSCs and M2 macrophages. The third pattern was associated with stromal components such as angiogenesis and extracellular matrix, as well mesenchymal cells and cancer stem cells. Furthermore, we analyzed the correlation between 61 TME-related signatures and the other TME-related signatures quantified by the MCP-counter algorithm in the CRC-AFFY cohort, with positive correlations of lymphocytic and stromal signatures with the signatures of the MCP-counter algorithm and highlighted the robustness of the different methods (**Fig. S1F**). Finally, we used the Kaplan-Meier method and Cox proportional hazard regression analysis to evaluate the prognosis of the TME-related signatures, and the stromal and tumor components significantly correlated with decreased survival (**Fig. S1G, Supplementary Table3-6**). Collectively, these data implied that the TME heterogeneity with distinct differences in immune, stromal, and metabolic reprogramming contributes to the development of tumors, and that the TME panel could be used to comprehensively characterize CRC.

### Determine and validation of CCCRC classification

With the increasing application of immunotherapy and tumor vaccines, there is growing evidence highlighting the importance of the TME in tumorigenesis and development (28, 29). To reveal the TME heterogeneity of CRC using the curated TME panel, consensus clustering analysis was performed based on the TME panel scores in the CRC-AFFY cohort, and the optimal cluster number was determined to be four using the consensus matrices heat map, CDF plot, and delta area plot (**Fig. S2A-C**). Subsequently, the CRC samples in the CRC-AFFY cohort were classified into the four CCCRCs with distinct TME components (**Fig. 1A-B, Fig. S2D**). PCOA showed that the four CCCRC subtypes were distinctly separated and the *P*-values for intercomparisons of the euclidean distances between them were all <0.05 using PERMANOVA test (**Fig. S2D**). The reproducibility of the CCCRC subtypes was externally validated in the CRC-RNAseq cohort and the same four CCCRC subtypes were revealed, with similar patterns of differences in the TME components (**Fig. S2E-G**). PCOA also demonstrated highly similar TME compartments in the same subtype between the CRC-RNAseq and CRC-AFFY cohorts (**Fig. S2D**). Differences in the TME components between the CCCRC subtypes were also observed in the analysis of previously reported immune and stromal signatures obtained by the MCP-counter, CIBERSORT, and ESTIMATE algorithm (**Fig. S3A-E**), and 10 classical oncogenic pathway activities and 86 metabolic pathway enrichment scores calculated by GSVA (**Supplementary Table7**).

**Figure 1.**
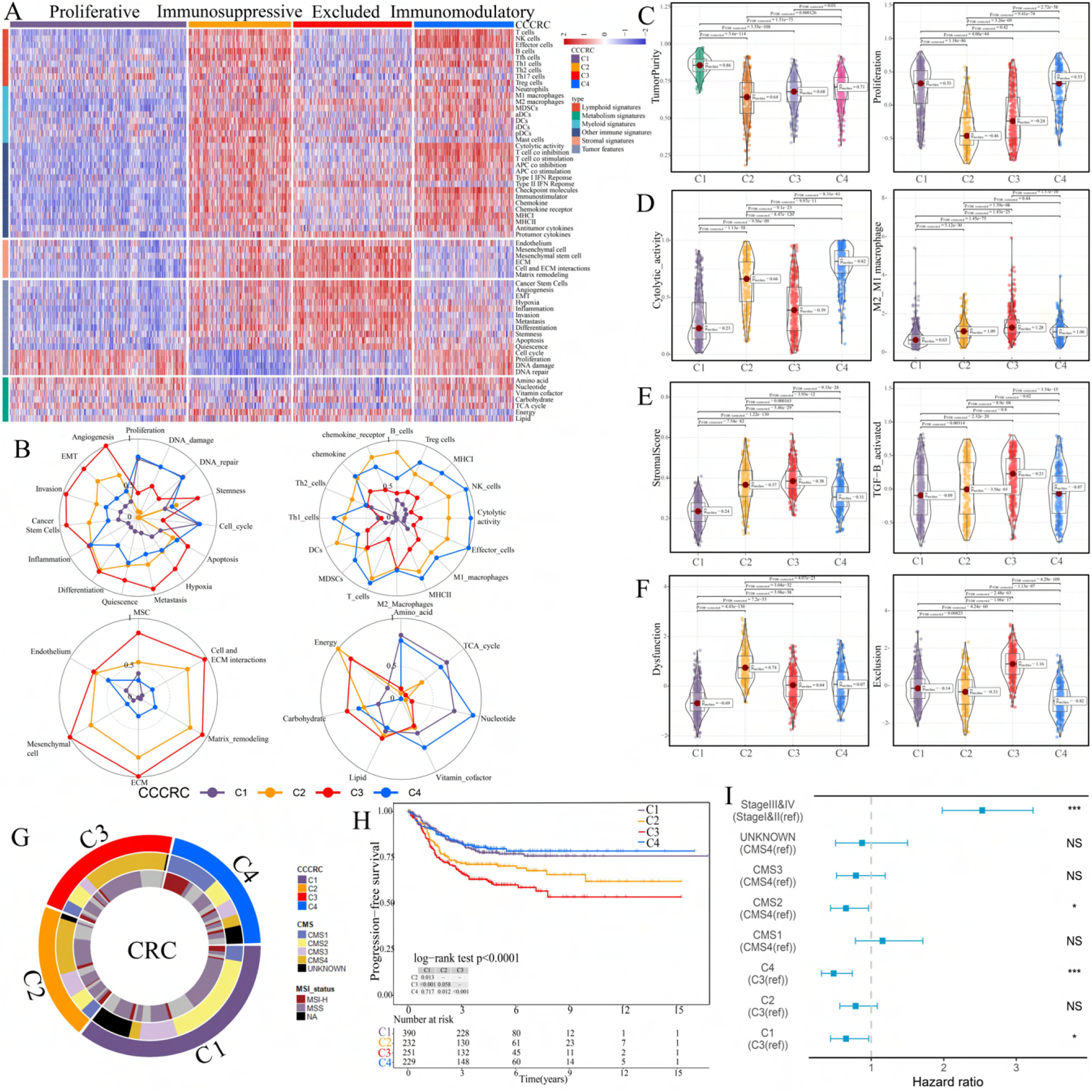
Comprehensive characterization of colorectal cancer (CCCRC). **A.** Heat map of 1471 CRC patients in the CRC-AFFY cohort classified into four distinct TME subtypes based on the 61 TME-related signatures. **B.** Radars display the characteristic TME-related signatures, including tumor, immune, stroma, and metabolism signatures, of each CCCRC subtype in the CRC-AFFY cohort. **C-E.** Box plots show differences in tumor (**C**), immune (**D**), and stroma (**E**) signatures in the CRC-AFFY cohort. Tumor purity and stroma scores were obtained from the ESTIMATE algorithm. Proliferative activity (proliferation), cytolytic score, M1 and M2 macrophage proportions, and TGFB activity were calculated by GSVA. **F.** Differences in T cell dysfunction and T cell exclusion scores between four CCCRC subtypes were analyzed based on the gene expression profiles in CRC-AFFY cohort. **G.** Overlap of CCCRC subtypes with consensus molecular subtypes (CMS) and microsatellite instability (MSI) status (high microsatellite instability [MSI-H], microsatellite stability [MSS]) in the CRC-AFFY and CRC-RNAseq cohorts. **H.** Kaplan-Meier method with log-rank test of progression-free survival (PFS) among the four CCCRC subtypes in the CRC-AFFY cohort. **I.** Forest plot of multivariate Cox proportional hazard regression analysis of PFS after adjusting for TNM stage and CMS subtype in the CRC-AFFY cohort. The hazard ratios are shown with 95% confidence intervals. *p value < 0.05; ***p value < 0.001; NS, p value > 0.05.

C1 (35% of all tumors), hereafter designated as the proliferative subtype, was characterized by the relative upregulation of tumor proliferative activity, tumor purity, and minimal or complete lack of lymphocyte and stromal infiltration, which was highly similar to the immune-desert phenotype previously described (**Fig. 1B-E**). The MYC, cell cycle and TP53 pathways associated with tumor proliferation had the highest GSVA scores in the C1 subtype (**Fig. S3E**). C2 (21% of all tumors), hereafter designated as the immunosuppressed subtype, was characterized by the relative upregulation of immune and stromal components, such as T cells, M2 macrophages, and cancer-associated fibroblasts (CAFs) (**Fig. 1B-E, Fig. S3A-D**). However, the extent of infiltration of effector cells, as well as the cytolytic score, was much lower than that of the C4 subtype (**Fig. 1B-E, E**). C3 (24% of all tumors), hereafter designated as the immune-excluded subtype, was characterized by the distinct upregulation of stromal components, such as CAFs, and cancer stem cells, as well as angiogenesis and hypoxia signatures (**Fig. 1B-E, Fig. S3A-D**). During tumor progression, TGF-beta secreted by CAFs is leveraged by tumor cells to suppress and exclude the anti-tumor immune components (30). We observed that the TGF-beta pathway, as well as WNT, NOTCH and RTK-RAS pathways, and the ratio of M2/M1 macrophages, were distinctly upregulated in C2 and C3 subtypes (**Fig. 1D-E, Fig. S3E**). The scores of 5/10 oncogenic pathways were the highest in the C3 subtype (**Fig. S3E**), suggesting that the activation of oncogenic pathways could lead to the formation of immune-excluded phenotypes which was consistent with the previous theory (31). C4 (20% of all tumors), hereafter designated as the immunomodulatory subtype, was characterized by the remarkable upregulation of anti-tumor-immune components, such as effector T cells, NK cells, and Th1 cells. The C4 subtype also had the highest cytolytic score compared with the other subtypes and lacked stromal components and the other immunosuppressed components, which indicated an immunomodulatory microenvironment (**Fig. 1B-E**).

To further explore the immune escape mechanism of each CCCRC subtype, the differences in T cell dysfunction and T cell exclusion scores between the four CCCRC subtypes were analyzed based on the gene expression profiles, which reflected the T cell features of the global tumor. Strikingly, the C2 subtype had highest T cell dysfunction score, indicating that T cell dysfunction in the C2 subtype was at the late stage (**Fig. 1F, Fig. S3F**). Using GSEA with all genes ranked according to the fold change between C2 and C4 subtypes, we found that terminally exhausted CD8+ T cell and TGF-beta signaling signatures were upregulated in the C2 subtype in the CRC-AFFY (**Fig. S3G**) and CRC-RNAseq (**Fig. S3H**) cohorts, which might reveal that CD8+ T cell infiltration within the tumor bed was suppressed by the stroma and was in a late state of dysfunction. The C3 subtype had the highest T cell exclusion score (**Fig. 1F, Fig. S3F**), demonstrating that the low T cell infiltration into the tumor bed was due to the increased abundance of CAFs, MDSCs, and M2 macrophages, thereby leading to the exclusion of T cells from the tumor bed.

Metabolic reprogramming also differed significantly among the four CCCRC subtypes (**Fig. 1B, Fig. S3I**). We analyzed the 86 metabolism pathways obtained from the KEGG database (**Supplementary Table7**) and observed that the number of upregulated metabolic pathways of the C3 subtype was the lowest. We also found that glycan metabolism was distinctly upregulated in C2 and C3 subtypes, which indicated that glycan metabolism was significantly associated with the stroma.

### Associations between CCCRC subtypes and other molecular subtypes and clinical characteristics

Previous studies have identified several molecular subtypes of CRC based on GEP. We investigated their associations with the CCCRC subtypes in the CRC-AFFY and CRC-RNAseq cohorts (**Fig. 1G, Fig. S4A-F**). The C1 subtype was primarily comprised of the CMS2 subtype and lower crypt-like subtype, and it contained the highest frequencies of the CCS1 subtype, B-type subtype, and TA subtype. The C2 subtype mainly consisted of the CMS4 subtype, surface crypt-like subtype, CCS3 subtype, C-type subtype, and inflammatory subtype, and included the highest frequency of the enterocyte subtype. The C3 subtype contained the highest frequencies of CMS4, CCS3, and C-type subtypes and was mainly comprised of the mesenchymal subtype and TA subtype. The C4 subtype included the highest frequencies of high microsatellite instability (MSI-H) and the CMS1 subtype, CIMP-H-like subtype, A-type subtype, and inflammatory subtype, and was mainly comprised of the CCS2 subtype.

We also focused on the differences in the TME components between the CCCRC subtypes and the CMS subtypes. Compared with the CMS1 subtype, the C4 subtype showed upregulated anti-tumor-immune components in the CRC-AFFY cohort and lacked immunosuppressive components, which were also found in the CRC-RNAseq cohort (**Fig. S5A**). CRC patients with MSI-H were sensitive to ICB treatment, with C4 and CMS1 subtypes containing approximately 47% and 75% of MSI-H, respectively. The C4 subtype with MSI-H showed upregulated scores of effector cells and cytolytic activity and downregulated scores of extracellular matrix and matrix remodeling compared with the CMS1 subtype with MSI-H (**Fig. S5B**). Similarly, we observed that the C4 subtype with MSI-H and the C4 subtype with MSS had higher scores of anti-tumor immune signatures and lower scores of stromal components, while the other CCCRC subtypes with MSI-H lacked anti-tumor immune signatures and had more stromal components (**Fig. S5C**). This analysis indicated that CCCRC subtypes could further classify the CMS subtype and MSI status to identify patients suitable for ICB therapy.

The Kaplan-Meier method showed that the C4 subtype had significantly higher overall survival (OS) and progression-free survival (PFS) than C2 and C3 subtypes, with the C3 subtype showing the worst OS and PFS (**Fig. 1H, Fig. S6A**). Multivariate Cox proportional hazard regression analyses also demonstrated that the C4 subtype independently predicted the best OS and PFS, whereas the C3 subtype independently predicted the worst OS and PFS after adjusting for TNM stage and CMS classification system (**Fig. 1I, Fig. S6B**). Similar results after the analysis of prognosis were observed in the CRC-RNAseq cohort (**Fig. S6C-F**).

### Differences in histological characteristics between CCCRC subtypes

To further explore the biological differences between CCCRC subtypes, we investigated the histological phenotypes by evaluating the WSIs of the TCGA-CRC cohort. We compared our CCCRC system with the three-category immune classification system of solid tumors, termed “desert”, “excluded”, and “inflamed” phenotypes (32, 33). Two pathologists evaluated the histological characteristics for each subtype under the microscope. The CRC samples in the TCGA-CRC cohort were categorized as these three phenotypes based on the abundance of lymphocytes and their spatial location with malignant epithelial cells. According to the three-category immune classification system, the C4 subtype was enriched with an inflamed phenotype characterized by abundant lymphocytes in direct contact with malignant cells (**Fig. 2A**). The C2 subtype was mostly categorized as an excluded phenotype. The C1 and C3 subtypes were mainly classified into the desert phenotype, whereas the C3 subtype was more frequently classified as an excluded phenotype than the C1 subtype. Notably, the lymphocytes of C2 subtype were more frequently intermixed with intra-tumor stromal components, whereas the lymphocytes of C3 subtype were more frequently excluded from the tumor bed and intermixed with adjacent-tumor stromal components, both of which were classified as excluded phenotype according to the three-category immune classification system.

**Figure 2.**
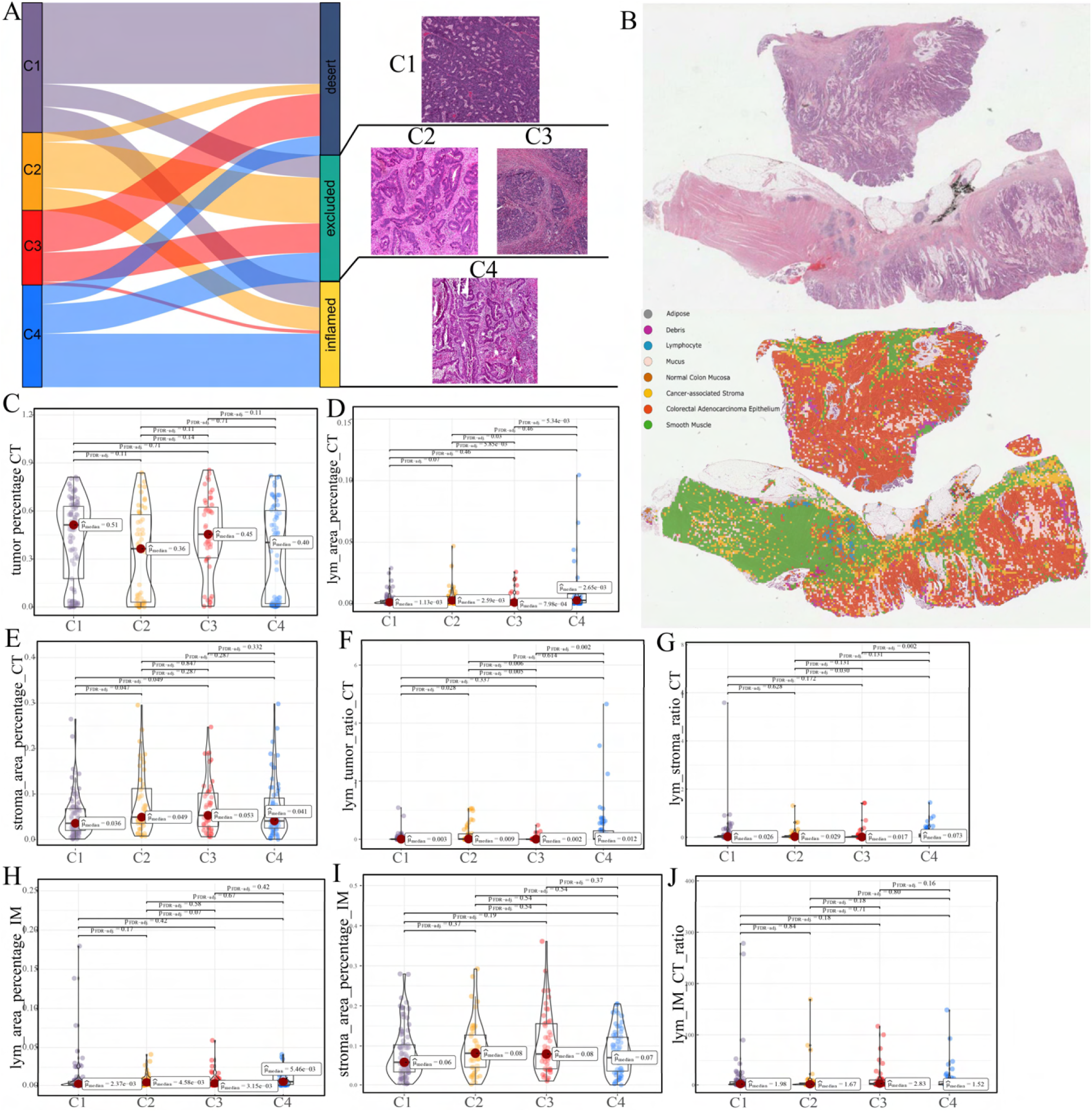
Differences in histological characteristics between CCCRC subtypes. **A.** Sankey plot shows overlap of CCCRC subtypes with the three-category immune classification system (“desert”, “excluded”, and “inflamed” phenotypes), and their representative hematoxylin and eosin (HE)-stained whole slide images (WSIs). C1: TCGA-AA-3955; C2: TCGA-A6-6654; C3: TCGA-CK-4948; and C4: TCGA-AD-6963. **B.** Representative WSI (top) and the CRC-multiclass model-inference segmentation of seven tissue types: tumor, stroma, lymphocyte, normal colon mucosa, debris, adipose, and mucin (bottom). **C-E.** Box plots show differences in the abundance of tumors (**C**), lymphocyte infiltration (**D**), and stroma (**E**) in the core tumor (CT) region. **F, G.** Box plots show differences in the lymphocyte infiltration to tumor content ratio (**F**) and lymphocyte infiltration to stromal content ratio (**G**) in the CT region. **H, I.** Box plots show differences in the abundance of lymphocytes infiltration (**H**) and stroma (**I**) in the invasive margin (IM) region. **J.** Box plots show differences in the ratio of lymphocyte infiltration in the IM region to the CT region.

The above differences in the histological characteristics among the CCCRC subtypes were based on the semi-quantitative analysis results of two pathologists, which are subjective to a certain extent. Therefore, we used hematoxylin and eosin (HE)-stained image-based deep learning to evaluate the abundance and spatial distribution of the tumor, lymphocytes, and stroma. The performance of our CRC-multiclass model was evaluated on the TCGA-CRC dataset with the accuracy reaching 81% and the AUCs for the different tissue types ranged from 0.95 to 0.98 (**Fig. S6G-H**). The tissue heatmap showed our model prediction results for a CRC WSI (**Fig. 2B**). In the core tumor (CT) region, the C1 subtype had a highly increased abundance of the tumor; the C4 subtype had increased lymphocyte infiltration and decreased stromal content; the C2 subtype had elevated lymphocyte and stromal infiltration; and the C3 subtype had the highest abundance of stroma, but less lymphocyte infiltration was detected (**Fig. 2C-E**). We also observed that C4 subtype had the highest lymphocyte infiltration to tumor content ratio and lymphocyte infiltration to stromal content ratio, followed by C2 subtype and C3 subtype had the lowest (**Fig. 2F, G**). In the invasive margin (IM) region, different degrees of lymphocyte infiltration and stromal components were observed for each subtype (**Fig. 2H, I**). Importantly, the ratio of lymphocyte infiltration in the IM region of the C3 subtype to the CT region was the highest, which confirmed that the stromal components excluded lymphocytes from the CT region in the C3 subtype (**Fig. 2J**). AI-enabled spatial analysis of WSIs confirmed the semi-quantitative results of the pathologists, with the C1 subtype belonging to the desert phenotype, C2 subtype belonging to the immunosuppressive phenotype, C3 subtype belonging to the excluded phenotype, and C4 subtype belonging to the hot phenotype. Collectively, our CCCRC system further refined the three-category immune classification system of solid tumors (32, 33) and conformed to the four-category immune classification system, termed “hot”, “desert”, “immune-excluded”, and “immunosuppressive” phenotypes (31).

### Biological characterization of CCCRC subtypes

We further elucidated the differences in biological characteristics among the CCCRC subtypes using multi-omics data from the TCGA and CPTAC databases, including genomics, epigenetics, transcriptomics, and proteomics data. As for the genomic alterations, the C4 subtype had the highest TMB and neoantigen values and the lowest prevalence of chromosomal instability (CIN), including SCNA counts and fraction of the genome altered (FGA) scores, compared with the other subtypes (**Fig. 3A, B**). Conversely, C1 and C3 subtypes displayed the highest CIN levels, as described by SCNA counts and FGA scores, and the lowest TMB and neoantigen values (**Fig. 3A, B**). The C2 subtypes displayed median CIN levels, TMB and neoantigen values. Among the frequently mutated genes (>5%), the mutation frequencies of *APC* (85.8%), *TP53* (64.9%), and *KRAS* (46.7%) were the highest in the C1 subtype compared to the other subtypes (all *P* < 0.05), followed by the C3, C2 and C4 subtypes, which are closely associated with the occurrence of CRC (**Fig. 3A, Supplementary Table8**). The C4 subtype was significantly enriched in mutations of *DNAH2* (26.0%), *MYH8* (26.8%), and *BRAF* (26.0%) genes (all *P* < 0.05), whereas the mutation frequency of C1, C2 and C3 subtypes was low. In terms of the differences in SCNA, the C1 subtype with the highest CIN level harbored significantly more amplified chromosomal regions (20q12, 20q13.12, 20q11.21, and 20q13.32) and deleted chromosomal regions (18q21.2, 18q22.1, and 18q12.3) (all *P* < 0.05) (**Fig. 3A, B, Supplementary Table9**). The C3 subtype was significantly enriched in the amplified chromosomal regions of 13q33.3, 13q22.1, and 13q12.2 and the deleted chromosomal regions of 8p21.2 and 8p23.2 (all, *P* < 0.05). No SCNA was significantly enriched in C2 and C4 subtypes. The single alteration events could not adequately delineate the CCCRC subtypes, we further computed the fraction of the altered samples per oncogenic pathway in each CCCRC subtype. The C4 subtype had the highest frequency of mutations in the cell cycle, HIPPO, MYC, NOTCH, PI3K, TGFB and RTK-RAS pathways (all *P* < 0.05) (**Fig. 3C, Supplementary Table10**). Notably, the C1 subtype had the highest frequency of mutations in the WNT pathway (*P* = 0.019). The frequency of mutations in the TP53 pathway was not significantly different between CCCRC subtypes. The 10 oncogenic pathways had higher frequencies of amplification (all *P* < 0.05), and 9 oncogenic pathways (except the NRF2 pathway) had higher frequencies of deletion (all *P* < 0.05) in C1 and C3 subtypes compared with C2 and C4 subtypes. Although none of genomic alterations was limited to or specific to a particular subtype, the apparent enrichment of certain alteration events within the CCCRC subtypes might highlight the TME heterogeneity and the genotype-CCCRC correlations of CRC.

**Figure 3.**
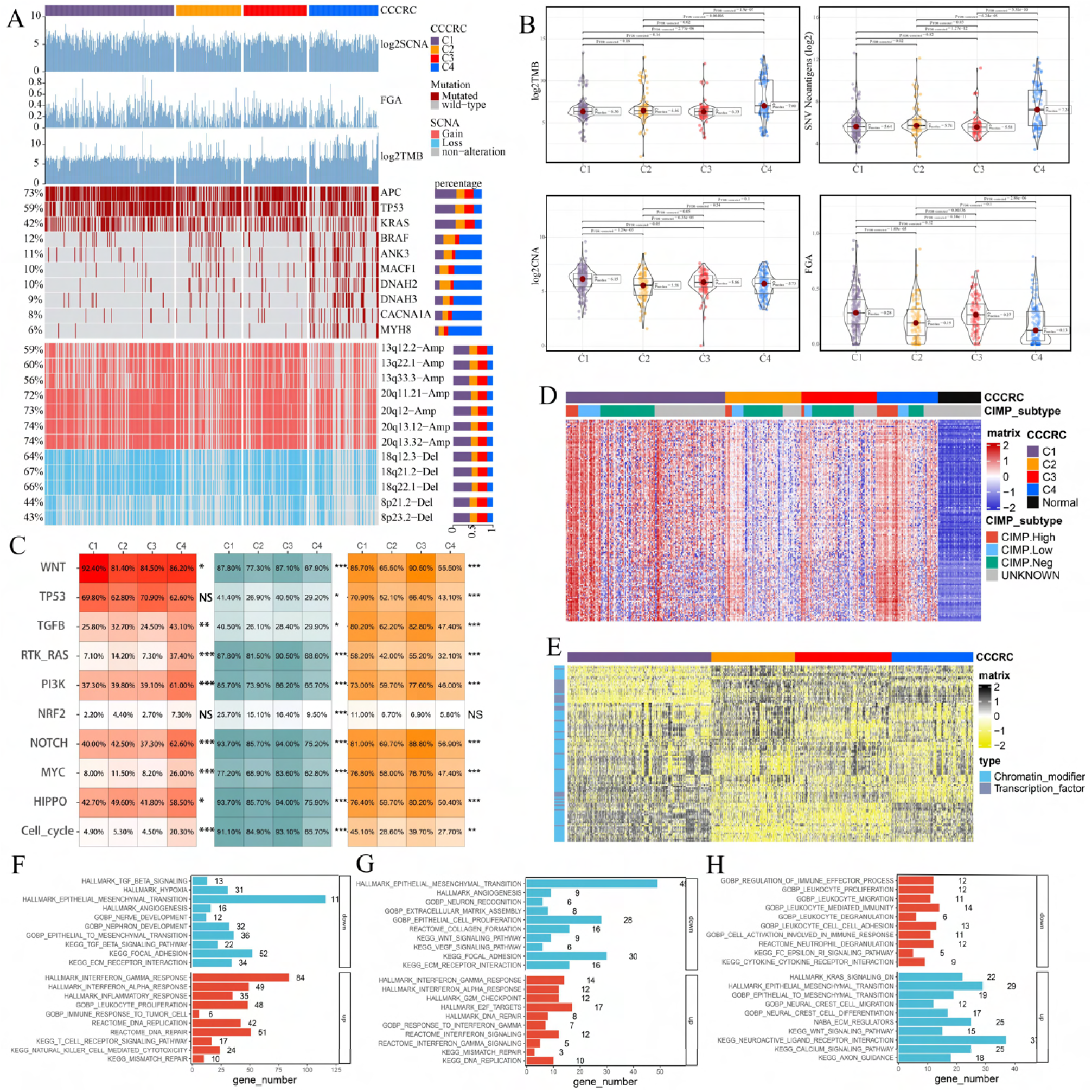
Biological characterization of CCCRC subtypes based on multi-omics data. **A.** Distribution of driver gene mutations and somatic copy number alterations (SCNAs) among the CCCRC subtypes in the TCGA-CRC dataset. **B.** Box plots show differences in tumor mutation burden (TMB), neoantigens, SCNA counts, and fraction of the genome altered (FGA) scores among the four CCCRC subtypes in the TCGA-CRC dataset. **C.** Genomic alterations in 10 oncogenic pathways were compared among the four CCCRC subtypes in the TCGA-CRC dataset. The color of the box represents the different types of genomic alterations (red, mutation; blue, amplification; yellow, deletion), and the color saturation represents the frequency. The color of the p value represents which oncogenic pathway had the highest frequency of the genomic alterations. **D, E.** Heat map shows differentially methylated genes derived from each CCCRC subtype vs normal tissues (**D**) and regulon activity profiles for transcription factors and chromatin modifiers (**E**). **F.** Significantly enriched gene sets among genes upregulated in the C4 subtype (red bars) and the C3 subtype (blue bars). **G.** Significantly enriched gene sets among proteins upregulated in the C4 subtype (red bars) and the C3 subtype (blue bars). **H.** Significantly enriched gene sets of methylated genes with downregulated DNA methylation in the C4 subtype compared to the C3 subtype (red bars) or with upregulated DNA methylation in the C4 subtype compared to the C3 subtype (blue bars). *p value < 0.05; **p value < 0.01; ***p value < 0.001; NS, p value > 0.05.

Subsequently, we found that the different CCCRC subtypes displayed highly diverse epigenetic, transcriptional, and proteomic profiles. As expected, the analysis of differentially methylated genes (DMGs) between CRC and normal tissues demonstrated that the C4 subtype had the most DMGs (n = 145) cared to the C1 subtype (n = 109), C2 subtype (n = 12), and C3 subtype (n = 23), and the C4 subtype exhibited extensive hypermethylation with the highest frequency of the CpG island methylator phenotype (CIMP) compared with the other subtypes (**Fig. 3D**). We further analyzed the regulon activity of critical chromatin modifiers and transcription factors in CRC, which could better evaluate their combinatorial biological effects. The regulon activity of the chromatin modifiers of the C1 subtype was generally higher than that of the other subtypes (**Fig. 3E**). The differences in the regulon activity of the chromatin modifiers might indicate that epigenetically driven transcriptional networks contributed to the remodeling of the TME, especially in the C1 subtype. Meanwhile, we observed that each subtype had different transcription factor activities (**Fig. 3E**). C1-specific upregulated genes (FDR < 0.001, top 1,000 by log_2_FC) were enriched for the pathways associated with tumor proliferation and metabolism (**Fig. S7A**). C2-specific upregulated genes were enriched for the pathways associated with immune function, stroma, and neurons (**Fig. S7A**). C3-specific upregulated genes were enriched for the pathways associated with stroma and neurons (**Fig. S7A**). Both C2 and C3 subtypes were enriched in neuron-associated pathways, suggesting that neuronal development might be involved in the formation of ECM (**Fig. S7A**). C4-specific upregulated genes were enriched for the pathways associated with anti-tumor immune function (**Fig. S7A**). The CCCRC-specific downregulated methylation genes (FDR < 0.001, top 1,000 by FDR) and the CCCRC-specific upregulated proteins (*P*-value < 0.05) were also enriched for analogous biological functional categories (**Fig. S7B, C**). Gene expression differences among the CCCRC subtypes were validated in the CRC-RNAseq cohort (**Fig. S7D-G**). DMGs, differentially expressed genes (DEGs), and differentially expressed proteins (DEPs) between each subtype were enriched for similar biological functional categories. Indeed, DEGs and DEPs upregulated in the C4 subtype compared with the C3 subtype were significantly enriched for immune-related pathways, whereas DEGs and DEPs upregulated in the C3 subtype compared with the C4 subtype were highly enriched for TGF beta signaling, EMT and angiogenesis (**Fig. 3F, G**). Similarly, genes with increased DNA methylation in the C4 subtype compared with the C3 subtype were enriched for EMT and ECM regulation, whereas genes with decreased DNA methylation in the C4 subtype were significantly enriched for immune-related pathways (**Fig. 3H**). Collectively, the similar differential biological patterns of DNA methylation, gene expression, and proteins among the CCCRC subtypes highlighted their role in influencing the TME of CRC.

### Discovery of a nongenetic tumor evolution pattern

Based on the theory of linear tumor evolution, we sought to investigate whether there is a dominant evolutionary pattern among the different CCCRC subtypes. We integrated DNA methylation, as well as transcriptomic and proteomic profiling, to analyze the differences between each pair of CCCRC subtypes. Strikingly, the evolutionary patterns from C1 to C4, C2, and C3 subtypes had the same sign in log2 (fold changes) and were dominate: all positive for increasing DNA methylation (FDR < 0.05) /gene expression (FDR < 0.05)/protein level (*P*-value < 0.05) or all negative for decreasing DNA methylation/gene expression/protein level (**Fig. 4A-C**). Furthermore, we intersected all the positives for increasing gene expression from C1 to C4, C2, and C3 subtypes in the CRC-AFFY and CRC-RNAseq cohorts and obtained 20 CCCRC genes (**Fig. 4D**, **Supplementary Table11**), which were associated with TGF-beta signaling and neural development. High expression of all 20 genes was significantly associated with poor PFS prognosis. To quantify the evolutionary pattern of individual CRC patients, we performed GSVA to generate CCCRC scores. To better evaluate the molecular features of the CCCRC scores, we also analyzed the correlation between the CCCRC scores and the TME panels. As expected, the CCCRC scores were strongly associated with the immunosuppressive signatures, including M2 macrophages, MDSCs, Treg cells, mesenchymal cells, EMT, angiogenesis, and hypoxia (**Fig. S7H**). The CCCRC score was the highest in the C3 subtype than in the other subtypes (**Fig. 4E**), and the high CCCRC score was significantly associated with shorter OS (**Fig. 4F**). Overall, our analysis implied that the four CCCRC subtypes not only had their own unique biological characteristics, but also had a dominant evolutionary pattern driven by epigenetic, transcriptional, and proteomic reprogramming.

**Figure 4.**
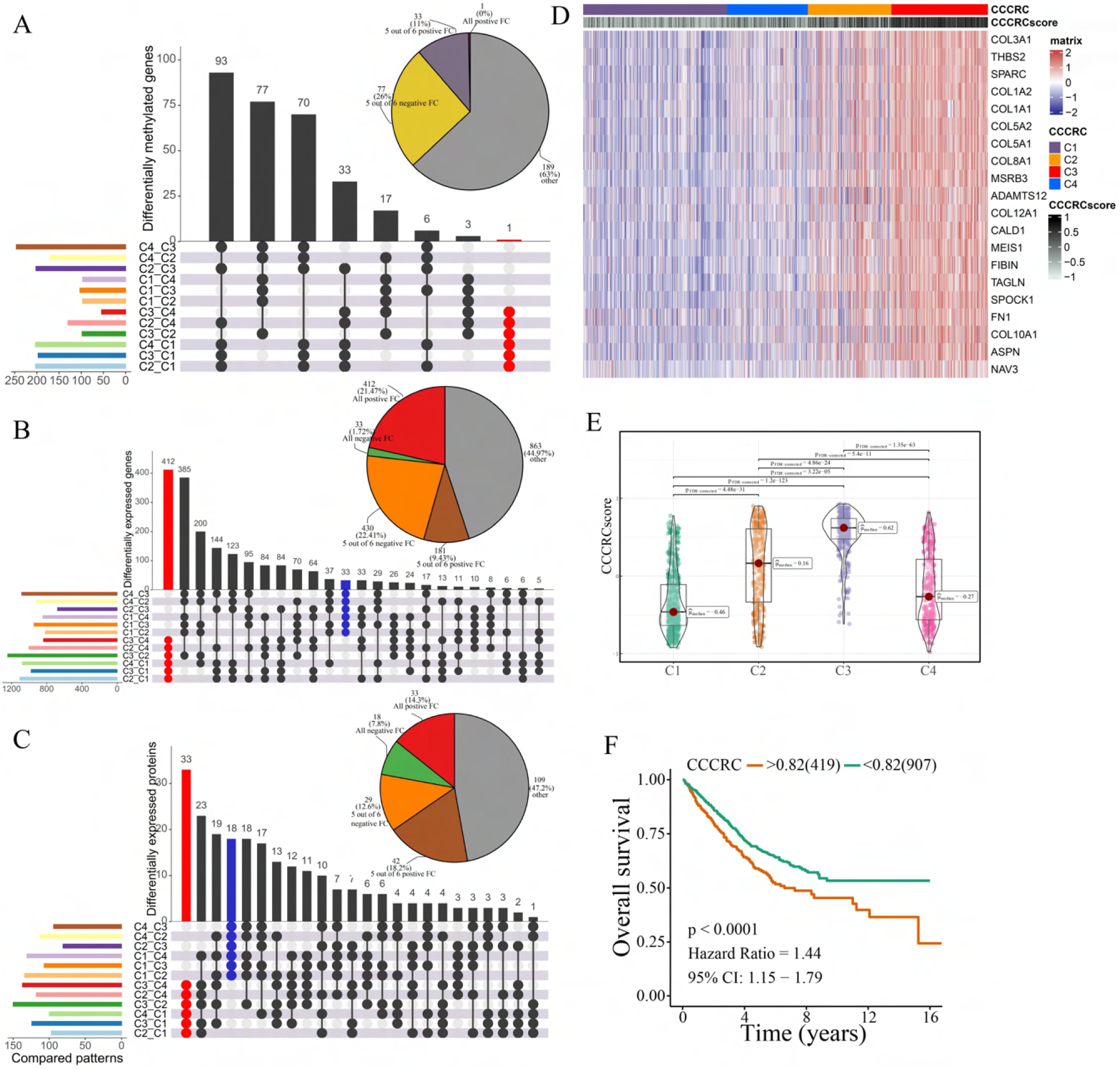
Discovery of a nongenetic tumor evolution pattern. **A-C.** Venn plots show “all positive” for increasing DNA methylation/gene expression/protein levels or “all negative” for decreasing DNA methylation/gene expression/protein levels. Pie chart (top right) distributions of the sign of pairwise FCs computed for differentially methylated genes (A), all differentially expressed genes (B) and all differentially expressed proteins (C). **D.** Heatmap shows gene expression levels of 20 CCCRC genes among the four CCCRC subtypes. **E.** Box plots show differences in the CCCRC score among the four CCCRC subtypes in the CRC-AFFY cohort. **F.** Kaplan–Meier method of overall survival (OS) among the four CCCRC subtypes in the CRC-AFFY and CRC-RNAseq cohorts.

### Differences in T cell function between CCCRC subtypes

We obtained the gene expression data for 7766 T cells from 12 patients with CRC, including four patients with the C1 subtype, one patient with the C2 subtype, two patients with the C3 subtype, and four patients with the C4 subtype (**Supplementary Table12**). A total of five CD4+ and four CD8+ T cell clusters were identified in tumor and normal tissues, including CD8+ intraepithelial lymphocytes (CD8+ IELs), effector memory CD8+ T cells (CD8+ Tem), recently activated effector memory or effector CD8+ T cells (CD8+ Temra/Teff), exhausted CD8+ T cells (CD8+ Tex), central memory CD4+ T cells (CD4+ Tcm) and naive CD4+ T cells (CD4+ Tn), tissue-resident memory CD4+ T (CD4+ Trm) cells, TH1-like cells, Treg cells, and T cycling cells (**Fig. S8A, B**). The characteristics of the T-cell clusters are summarized in **Supplementary Table13**. **Fig. 5A** and **B** show the distribution of the 10 T cell clusters among each CCCRC subtype. The bulk RNAseq analyses demonstrated that C2 and C4 subtypes showed relative upregulation of immune components. Notably, we found that the C4 subtype was enriched in CD8+ Tem and CD8+ Temra/Teff cells, but lacked CD8+ Tex cells compared with the C2 subtype (**Fig. 5C, D**). Within the subset of CD8+ Tex cells, we distinguished two smaller subsets according to their gene expression markers, KLRG1+ CD8+ Tex cells and HSPA1B+ CD8+ Tex cells (**Fig. S8C, D**). KLRG1+ CD8+ Tex cells were more enriched in C2 and C3 subtypes than the C4 subtype (**Fig. 5E**), which resemble terminally exhausted T cells, and they were associated with non-response to ICB therapy (34). Moreover, the higher ratio of KLRG1-to-CD8A expression, the worse the OS of patients in CRC-AFFY and CRC-RNAseq cohorts (**Fig. 5F, G**). Meanwhile, we re-clustered the Treg cells and identified four Treg cell subsets, namely, TXNIP+ Treg cells, TNFRSF4+ Treg cells, HSPA1A+ Treg cells, and IFIT1+ Treg cells (**Fig. S8E-H**). We found that TNFRSF4+ Treg cells were significantly more enriched in C2 and C3 subtypes than the C4 subtype (**Fig. 5H**), which might indicate that TNFRSF4+ Treg cells were closely related to the formation of the tumor stroma. The higher ratio of TNFRSF4-to-FOXP3 expression, the worse the OS of patients in CRC-AFFY and CRC-RNAseq cohorts (**Fig. 5I, J**). Equally important, patients with a high ratio of KLRG1-to-CD8A expression or a high ratio of TNFRSF4-to-FOXP3 expression who received ICB therapy had a shorter OS and PFS than those with a low ratio of KLRG1-to-CD8A expression or a low ratio of TNFRSF4-to-FOXP3 expression in Gide, Hugo, Jung, and IMvigor210 datasets (**Fig. S9A-H)**. We also found that the expression of KLRG1 and TNFRSF4 was higher in CD8+ T cells and Treg cells, respectively, in tumor tissues than in adjacent tissues (**Fig. 5K, L**). Overall, we used scRNAseq data to analyze the differences in T cell function among the different CCCRC subtypes, and the C2 subtype did show more immunosuppression than the C4 subtype, which was consistent with the bulk RNAseq analyses.

**Figure 5.**
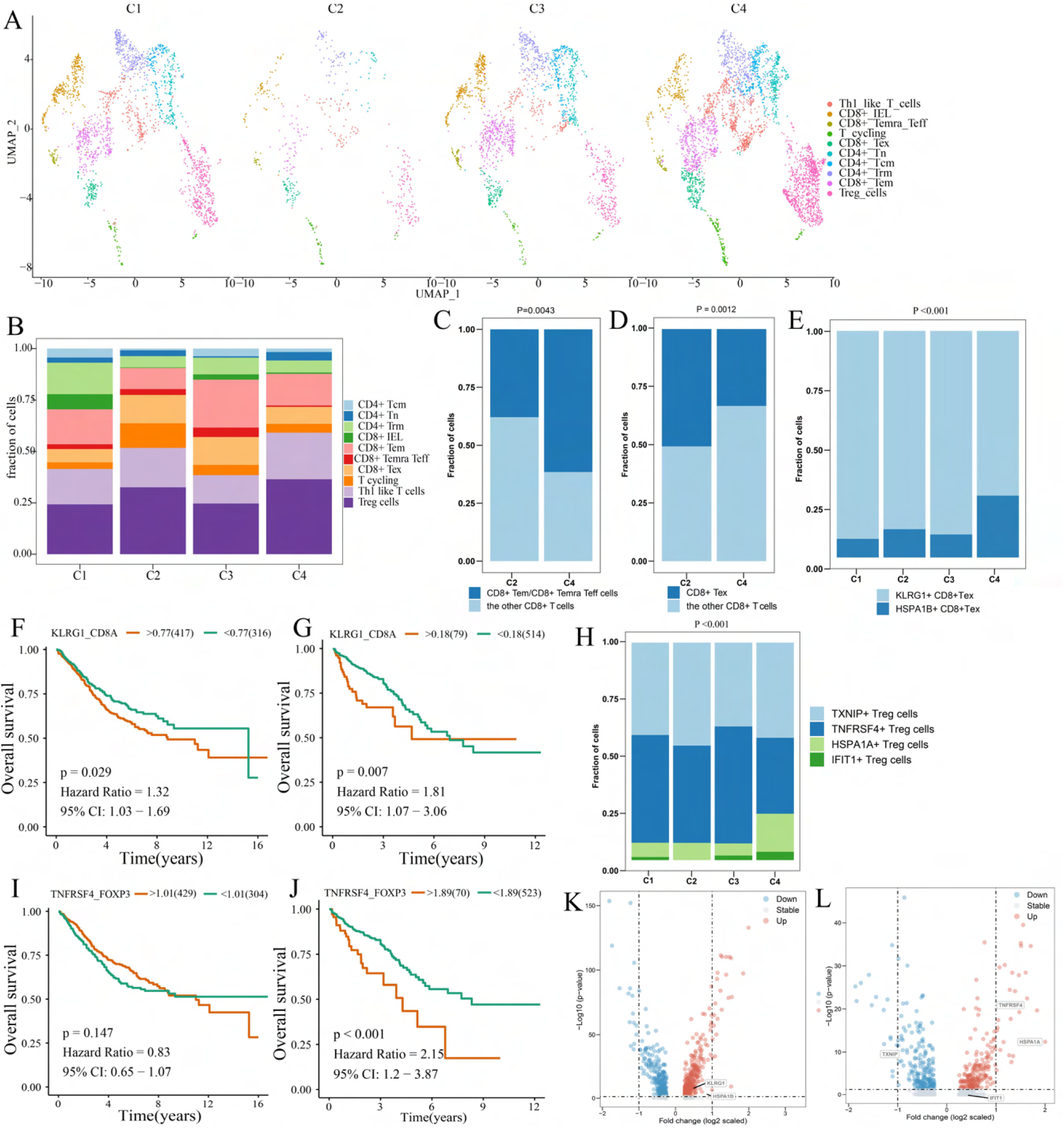
Differences in T cell function between CCCRC subtypes. **A.** UMAP shows the composition of T cells colored by cluster and divided by the CCCRC subtype in CRC tissues. **B.** Histogram shows the cell distribution of 10 T cell types in the different CCCRC subtypes. **C.** Proportion of effector memory CD8+ T cells (CD8+ Tem), recently activated effector memory or effector CD8+ T cells (CD8+ Temra/Teff), and the other CD8+ T cells (shown in the histogram) in the C2 and C4 subtypes. **D.** Proportion of exhausted CD8+ T cells (CD8+ Tex) and the other CD8+ T cells (shown in the histogram) in the C2 and C4 subtypes. **E.** Histogram shows the cell distribution of KLRG1+ CD8+Tex and HSPA1B+ CD8+Tex cells in the different CCCRC subtypes. **F, G.** Kaplan-Meier method with log-rank test of overall survival (OS) in the CRC-AFFY cohort (**F**) and the CRC-RNAseq cohort (**G**) between low and high ratios of KLRG1-to-CD8A expression in patients. **H.** Histogram shows the cell distribution of TXNIP+ Treg cells, TNFRSF4+ Treg cells, HSPA1A+ Treg cells, and IFIT1+ Treg cells in the different CCCRC subtypes. **I, J.** Kaplan–Meier method with log-rank test of OS in the CRC-AFFY cohort (**I**) and the CRC-RNAseq cohort (**J**) between low and high ratios of TNFRSF4-to-CD8A expression in patients. **K.** Volcano plot shows differentially expressed genes between tumor (red dots) and normal CD8+ T cells (blue dots). **L.** Volcano plot shows differentially expressed genes between tumor (red dots) and normal Treg cells (blue dots).

### **S**ignificance of CCCRC in guiding clinical treatment of CRC

The 5-fluorouracil (5-FU)-based chemotherapy, anti-VEGF (bevacizumab), and anti-EGFR (cetuximab, panitumumab) therapies are the first-line treatment options for CRC. We further explored whether the different CCCRC subtypes could predict therapeutic efficacy. In the CRC-AFFY cohort, 564 patients with stage II and III CRC had chemotherapy-related clinical information, including 323 who were not treated by chemotherapy and 241 who were treated by chemotherapy. Furthermore, 155 stage II and III CRC patients with or without chemotherapy in the GSE103479 dataset were also included in our study. We found that C1 patients with stage II and III CRC receiving chemotherapy had a better OS than those who did not and were more suitable for 5-FU-based chemotherapy in the CRC-AFFY cohort and the GSE103479 dataset (**Fig. 6A, B**).

**Figure 6.**
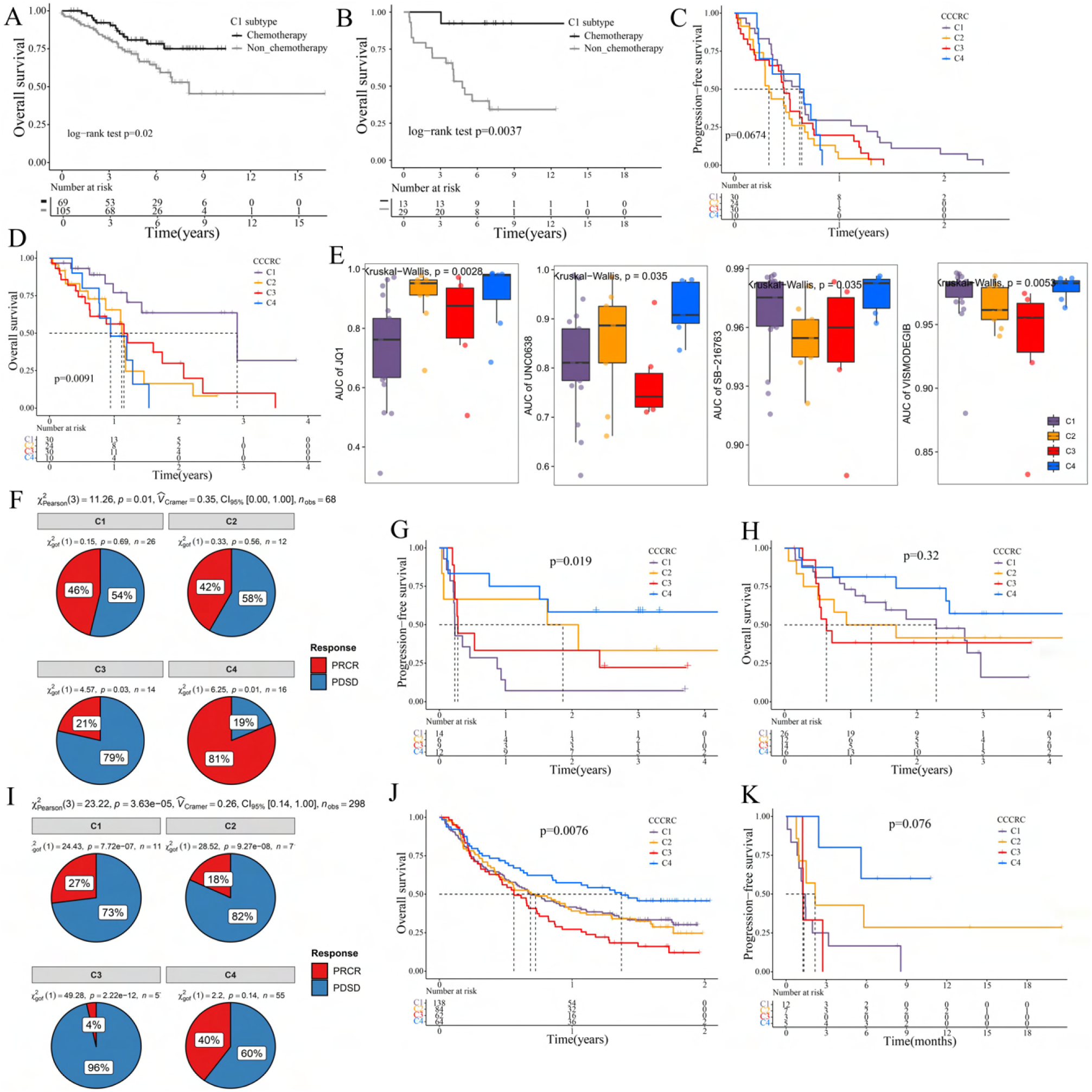
Significance of CCCRC in guiding the clinical treatment of colorectal cancer. **A, B.** Kaplan-Meier method of overall survival (OS) between stage II and III CRC C1 patients with or without chemotherapy in the CRC-AFFY cohort (**A**) and the GSE103479 (**B**) dataset. **C, D.** Kaplan-Meier method of OS (**C**) and progression-free survival (PFS) (**D**) among the four CCCRC subtypes in the GSE104645 dataset. **E.** Box plots show the differences in the area under the receiver operator characteristics curve (AUC) of drug responses among the four CCCRC subtypes. **F.** Pie chart shows the differences in the proportion of responses to immune checkpoint blockade treatment among the four CCCRC subtypes in the two independent melanoma cohorts (Gide and Hugo datasets, n = 68) treated with anti-PD1 therapy. **G, H.** Kaplan-Meier method with log-rank test of PFS (**G**) and OS (**H**) among the four CCCRC subtypes in the two independent melanoma cohorts (Gide and Hugo datasets, n = 68) treated with anti-PD1 therapy. **I.** Pie chart shows the differences in the proportion of responses to immune checkpoint blockade treatment among the four CCCRC subtypes in the urothelial carcinoma cohort (n = 298) treated with anti-PDL1 therapy. **J, K.** Kaplan-Meier method with log-rank test of OS and PFS among the four CCCRC subtypes in the urothelial carcinoma cohort (n = 348) (**J**) and the lung cancer cohort (n = 27) (**K**) treated with anti-PD1/PDL1 therapy. PRCR: partial response and complete response; PDSD: progressive disease and stable disease.

Furthermore, 162 mCRC patients were treated with chemotherapy or a combination of chemotherapy and bevacizumab in the GSE104645 dataset. The response rate (RR) after chemotherapy (including partial response [PR] and complete response [CR]) of C1 and C4 subtypes tended to be higher than that of C2 and C3 subtypes (**Fig. S10A**), whereas the RR of the C2 subtype treated with a combination of chemotherapy and bevacizumab tended to be higher than that of the other subtypes (**Fig. S10B, C**). In addition, the RR tended to be higher in the C2 subtype treated with (5-FU)-based chemotherapy and bevacizumab than in those treated with chemotherapy alone (**Fig. S10D**).

The GSE104645 dataset also contained 111 mCRC patients without the *RAS* mutation who were treated with anti-EGFR antibody. The disease control rates (DCR) after anti-EGFR therapy (including partial response, complete response, and stable disease) were 75% for C1, 66% for C2, 51% for C3, and 65% for C4, respectively (*P* = 0.16) (**Fig. S10E**). The DCR of the C1 subtype with anti-EGFR therapy tended to be higher than that of the other subtypes (*P* = 0.08) (**Fig. S10F**). Notably, PFS of the C1 subtype with anti-EGFR therapy tended to be better than that of the other subtypes (log-rank *P*-value = 0.067) and OS of the C1 subtype was significantly better than that of the other subtypes (log-rank *P*-value = 0.0091) (**Fig. 6C, D**). The above results suggested that the C1 subtype may benefit from chemotherapy and anti-EGFR treatment, whereas the C2 subtype may benefit from a combination of (5-FU)-based chemotherapy and bevacizumab, but there was no evidence that the C3 subtype is suitable for these treatments.

To further explore the treatment strategies of the CCCRC subtypes, we trained a pre-clinical model based on a filtered gene set comprised of 81 CCCRC subtype-specific and cancer cell-intrinsic gene markers (**Supplementary Table14**). The pre-clinical model was constructed using the xgboost algorithm with the highest accuracy, AUC and F1 scores (**Fig. S11A-C)**. The 71 human CRC cell lines were classified into four CCCRC subtypes (**Supplementary Table15**). The AUCs of the drug response between CCCRC subtypes were compared (**Fig. 6E**). Notably, the AUCs of the bromodomain and extra-terminal domain inhibitor (BET) JQ1 was significantly lower in C1 subtype. The AUCs of G9a-specific inhibitor UNC0638 were significantly lower in the C3 and C1 subtypes. The AUCs of WNT pathway inhibitor SB216763 and Hedgehog pathway inhibitor vismodegib were significantly lower in the C3 and C2 subtypes.

Immune checkpoint blockade (ICB) therapy has recently emerged as a highly promising therapeutic strategy for various malignancies, but it lacks effective markers to identify suitable patients. We collected multiple ICB therapy-associated datasets to evaluate whether the CCCRC classification system could be used as a tool to predict ICB therapy efficacy. GSVA of the TME-related signatures and the Z-score normalization of signature scores could reduce the tissue-type-specific effects. In two independent melanoma datasets (Gide and Hugo datasets, n = 68) treated with anti-PD1 therapy, patients were classified into the four CCCRC subtypes. As expected, the RR to anti-PD1 therapy in the C4 subtype was 81% in contrast to only 21% in the C3 subtype (**Fig. 6F**), with prolonged PFS and OS in both subtypes (**Fig. 6G, H**). Similar findings were observed in the cohorts of anti-PD1/PDL1 treated patients with urothelial carcinoma (IMvigor210 dataset, n = 348) and lung cancer (Jung dataset, n = 27). RR was significantly higher in patients with the C4 subtype (40%) compared with the other subtypes (C1 with 17%, C2 with 18%, C3 with 4%) in the IMvigor210 dataset (**Fig. 6I**). The C1 subtype in the IMvigor210 and Jung datasets had the longest OS, while patients with the C3 subtype had the worst OS (**Fig. 6J, K**).

### Single-sample gene classifier construction

For each subtype, we selected the genes with FDR <0.05 and logFC >0 and ordered them according to fold-change to generate a subtype-specific gene set (n = 9,256 mRNA genes). After screening by the Boruta importance test, a total of 80 unique genes were used to construct the final classifier in the training set and the validation set (**Supplementary Table16**). As shown in **Fig. S11D-F**, the performance of the xgboost algorithm was the best with the highest accuracy, AUC and F1 scores. The gene classifier based on the xgboost algorithm is publicly available at https://github.com/XiangkunWu/CCCRC, and the CCCRC subtype information of a single patient can be obtained by directly inputting the gene expression matrix of the patient. The single-sample gene classifier could facilitate the discovery of new biomarkers and the personalized treatment of clinical patients with CRC.

## Discussion

The key role of the TME in dynamically regulating tumor progression and affecting treatment outcomes has been widely recognized, and treatment strategies targeting the TME have become a promising approach for cancer therapy (28,35–37). However, there are few comprehensive analyses that consider the tumor cells and the TME as a whole. The comprehensive dissection of the crosstalk between tumor cells and TME may reveal new tumor biology concepts and identify therapeutic targets, and ultimately achieve precise medical treatment (20, 28). Thus, we collected the molecular features of the tumor cells and TME to reconstruct the whole tumor composition and performed integrated analyses to understand the TME. The four CCCRC subtypes had distinct molecular and histopathological characteristics, therapeutic efficacy, and prognosis (**Fig. 7**). We identified a nongenetic evolutionary pattern from C1, C4, C2, and C3 was associated with an evolution from a cold (C1) to a hot (C4) and eventually suppressive (C2) and excluded (C3) microenvironment (**Fig. 7**).

**Figure 7.**
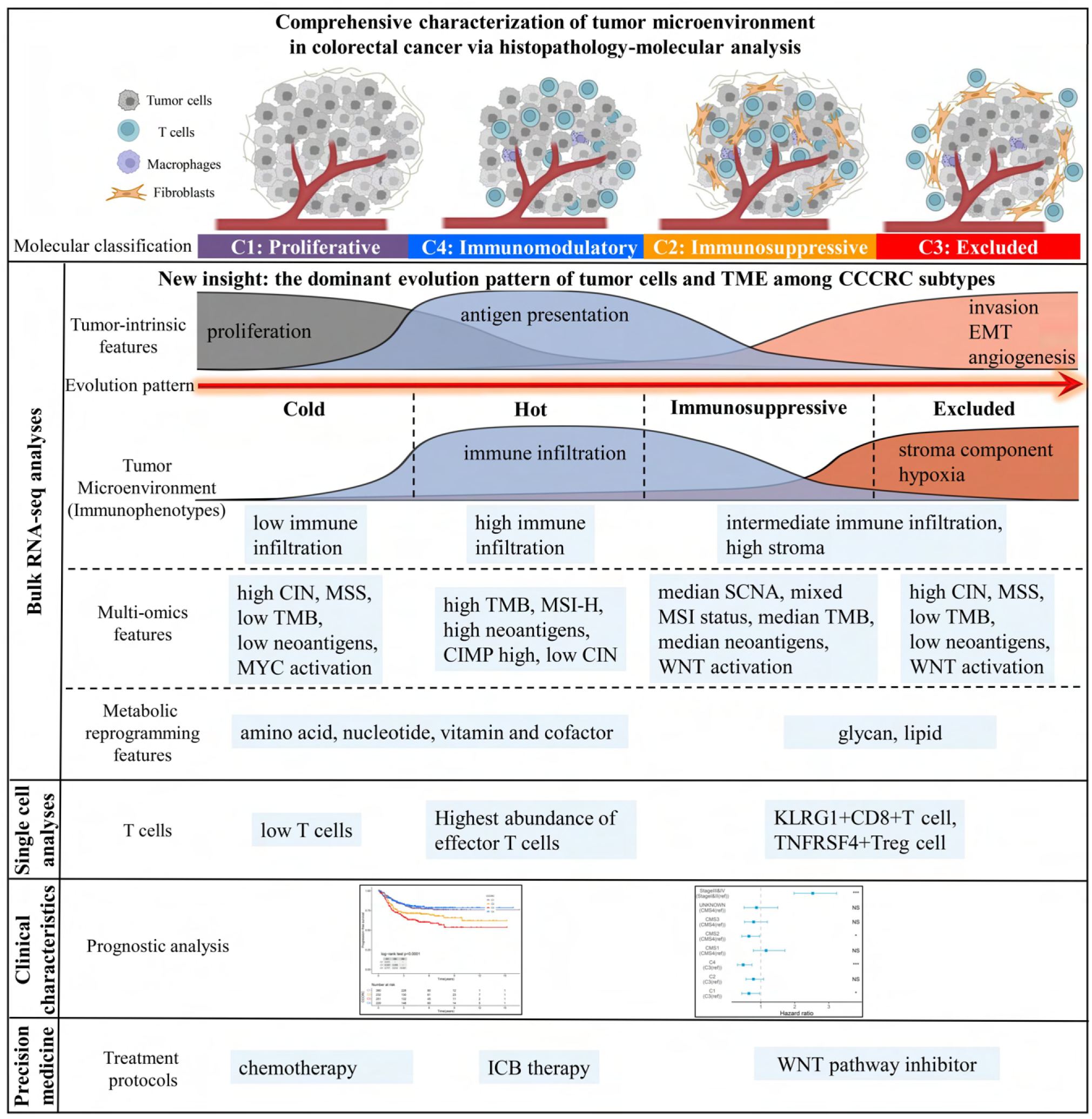
Overview of characteristics of CCCRC subtypes. These included tumor microenvironment features, multi-omics features, scRNA-seq features, treatment strategies and prognostic value for CCCRC subtypes.

In this study, we identified four subtypes with distinct TME features through unsupervised clustering analysis of approximately 2,000 CRC patients. C1 and C4 subtypes are typical desert and inflamed tumors, respectively, while C2 and C3 subtypes were difficult to classify into one of the classical immunophenotypes of the three-category immune classification system (“desert”, “excluded”, and “inflamed” phenotypes) (32, 33) based on TME features due to the unclear distribution of stromal components and lymphocytes. Our pathologists evaluated the histological characteristics for each subtype under the microscope and observed that the C2 subtype was mainly categorized as an excluded phenotype and the C3 subtype was mainly classified as a desert and an excluded phenotype. However, the WSIs showed that lymphocytes in the C2 subtype were more frequently intermixed with the stroma within but not adjacent to the main tumor mass, and lymphocytes in the C3 subtype were more frequently excluded from the tumor mass but not intermixed with lymphocytes within the main tumor mass, both of which were classified as the excluded phenotype. Notably, we used AI-enabled spatial analysis of WSIs to confirm the semi-quantitative results of the pathologist, that is, the C2 subtype had increased lymphocyte and stromal infiltration in CT and IM regions and the C3 subtype had the highest abundance of stroma and less lymphocyte infiltration in the CT region, while lymphocyte infiltration and stromal components were observed in the IM region. We also found that the C2 subtype had the highest T cell dysfunction score and the C3 subtype had the highest T cell exclusion score. GSEA demonstrated that the terminally exhausted CD8+ T cell signature was upregulated in the C2 subtype compared with the C4 subtype. scRNA-seq analysis showed that KLRG1+ CD8+ T cells were significantly more enriched in C2 and C3 subtypes than the C4 subtype. KLRG1+ CD8+ T cells were associated with nonresponse to ICB therapy, which were more terminally differentiated than KLRG1− CD8+ T cells and had lower proliferative capacity (34). KLRG1 is a marker of terminal differentiation of CD8+T cells (34), and the inhibitory receptor of ILC1s (group 1 innate lymphoid cells), ILC2s, and NK cells (38). ILC1s in tumors express high levels of the KLRG1 gene and pro-angiogenic activity and may even promote tumor progression in TGF-beta-rich tumors (38). Therefore, we defined C2 and C3 subtypes as immunosuppressed and immune-excluded, respectively. Our CCCRC classification system refined the three-category immune classification system (32, 33). Moreover, we defined for the first time the four-category immune classification system based on multi-omics analysis and histological characteristics (“hot”, “immunosuppressed”, “excluded”, and “cold” phenotypes) (31).

Interestingly, we observed a dominant evolution pattern among the CCCRC subtypes based on the theory of linear tumor evolution (39), that is, the evolutionary pattern from C1 (proliferative subtype) to C4 (immunomodulatory subtype), C2 (immunosuppressed subtype), and C3 (immune-excluded subtype) subtypes. We hypothesized that during the development of CRC, immune infiltration gradually increased with the increase of genomic alterations and tumor immunogenicity, while the stroma and nerves also gradually increased. The stroma and nerves play important roles in the progression of CRC, gradually causing lymphocytes to become exhausted and excluding them from the tumor bed. Tavernari et al. demonstrated that progression from lepidic to solid histology of lung adenocarcinoma was associated with a transition from a cold (lepidic) to a hot (papillary and especially acinar) and eventually suppressive and excluded (solid) microenvironment (40). Their proposed nongenetic tumor evolution pattern is consistent with our findings in CRC. What’s more, we have identified a gene list that promotes this evolutionary pattern and interfering with these genes may prevent tumor progression. We proposed CCCRC score based on the gene list to quantify the evolutionary pattern of individual CRC patients, which were independent prognosis predictors and associated with immunosuppressive components. Additional experimental evidence is needed to verify the bold speculation of this evolutionary pattern, and a large collective effort is needed to arrive at a consensus.

The CCCRC subtypes significantly correlated with previous molecular subtypes, including CMS subtypes (13), Budinska subtypes (6), De Sousa subtypes (7), Roepman subtypes (9), and Sadanandam subtypes (10), as well as prognosis. The CMS classification system integrates six independent classification systems utilizing a network-based approach (13), which is considered as the most robust classification system that is used to predict prognoses and to guide ICB therapy, chemotherapy, and anti-EGFR therapy as well as to screen new potential targeted drugs (41–46). However, patients with the CMS1 subtype, characterized by immune infiltration and activation, did not have the best prognoses compared with the other CMS subtypes, while patients with the CMS2 subtype, characterized by low immune and inflammatory signatures, had the best prognoses (13, 47). Our CCCRC subtypes significantly correlated with OS and PFS of patients and had higher correlation rates compared with the CMS classification system. We found that the CMS1 subtype showed fewer anti-tumor immune components and more stromal components and other immunosuppressive components compared to the C4 subtype. Meanwhile, the C4 subtype with MSI-H had higher immune infiltration compared with the CMS4 subtype with MSI-H. Thus, we boldly speculated that our CCCRC classification system was more suitable than the CMS classification system for predicting the prognosis and efficacy of ICB therapy.

The CCCRC classification system might facilitate clinical treatment decisions and new therapeutic target discoveries. To explore the potential treatment strategies for the CCCRC subtypes, we generated a gene list comprised of subtype-specific, cancer cell-intrinsic genes according to the study of Peter et al. to develop a pre-clinical model (41), which could be used to analyze the drug response data from cell lines, patient-derived xenografts, and tumor organoids. We observed that the C1 and C3 subtypes had higher CIN level than C2 and C4 subtypes. And most of the critical chromatin modifications had higher regulon activity in the C1 subtype. It has been well established that CIN and epigenetic silencing leads to decreased tumor intrinsic immunogenicity (48–50). Our analysis also demonstrated that the C1 subtype was more sensitive to the BET inhibitor JQ1. Zhang et al. found that JQ1 induces anti-tumor immunity in head and neck squamous cell carcinoma by enhancing MHC class I expression and can improve the response rate to ICB treatment (51). C1 and C3 subtypes were suitable for G9a-specific inhibitor UNC0638. Zhang et al. also found that BRD4 inhibits the MHC class I expression by recruiting G9a (51). The C2 and C3 subtypes were significantly enriched in the WNT pathway, and our analysis also indicated that these two subtypes were more sensitive to the WNT pathway inhibitor SB216763. Meanwhile, we identified a large number of mutant genes significantly enriched in the C4 subtype, which mutated to cause substantial immune infiltration and could be candidate genes for mRNA vaccine development. The RNA-mediated immunotherapy regulating the TME is known as the next era of cancer treatment (36). The CCCRC subtype-specific genes were also identified in our study to screen out the new therapeutic targets for the TME.

To conclude, our study proposed the CCCRC classification system and performed integrated data analysis to clearly characterize the molecular features and histological characteristics of each CCCRC subtype, develop the corresponding personalized treatments for patients with the different CCCRC subtypes, and construct the simple gene classifier to facilitate clinical application. We believe that our study will serve as a research paradigm for dissecting the TME and for transitioning from molecular classification to clinical translation, thereby accelerating the understanding of the TME in CRC and contributing to the development of therapeutic targets against TME.

## Supplementary material and methods

### Acquisition and processing of gene expression profiles (GEP) for the investigation of CCCRC

A total of 2195 samples were obtained from ten publicly available datasets (**Supplementary Table1**). The eight publicly available raw microarray datasets sequenced by the Affymetrix gene chip were downloaded from the Gene Expression Omnibus (GEO; https://www.ncbi.nlm.nih.gov/geo/) and renormalized by the robust multi-array average (RMA) method, including GSE13067, GSE13294, GSE14333, GSE17536, GSE33113, GSE37892, GSE38832 and GSE39582. Samples that overlapped in GSE14333 and GSE17536 datasets were excluded from the GSE14333 dataset. Level-3 TCGA and CPTAC RNA sequencing (RNAseq) datasets were obtained from the TCGA data portal (March 2022) (https://portal.gdc.cancer.gov/), and the count data were normalized by the “voom” method. Ensembl IDs were annotated into gene symbols using GENCODE (v36). If the gene symbol had multiple probes or duplicates, the median value was calculated as its relative GEP. Before merging the microarray datasets or RNAseq datasets into the CRC-AFFY or CRC-RNAseq cohort, the batch effects were examined using principal component analysis (PCA) and corrected using the “Combat” function. The selection criteria of these patients included: (1) CRC primary tissue samples; (2) coming from the same sequencing platform; (3) surgically resected specimens. The exclusion criteria included: (1) CRC metastatic tissue; (2) puncture tissues. Detailed information on the sample size and the corresponding clinicopathological data of the CRC-AFFY and CRC-RNAseq cohorts are summarized in **Supplementary Table1**.

### Calculation of the TME-related signature scores

After reviewing previously published studies, the Molecular Signatures Database (MSigDB; http://www.gsea-msigdb.org/gsea/msigdb/index.jsp), and the Reactome pathway portal (https://reactome.org/PathwayBrowser/), we identified relevant biomarker genes for tumor, immune, stromal, and metabolic reprogramming signatures. The 4,525 mRNAs from each of the 61 TME-related signatures are listed in **Supplementary Table2**, as well as the source of each signature. Gene set variation analysis (GSVA) with default parameters using R package “GSVA” was performed to calculate the signature score of each TME-related signature for each sample of each cohort separately based on the relative GEP (52).

### Normal tissue versus tumor tissue analysis

To assess the distribution of normal and tumor samples in the GSE39582 (n = 19 normal) and TCGA (n = 41 normal) datasets, the gene expression data of each dataset were re-normalized, including the normal samples (consistent with the description of data normalization above). Principal coordinate analysis (PCOA) based on euclidean distance was used to analyze the distribution between normal and CRC samples (53). Permutational multivariate analysis of variance (PERMANOVA) test was used to evaluate whether the difference in euclidean distances between the normal and CRC samples was statistically significant (obtained using R package “vegan” (54)).

### Comprehensive characterization of CRC

The “ConsensusClusterPlus” function of the R package “ConsensusClusterPlus” (26) was applied to identify the optimal number of CCCRC based on the TME-related signatures in the CRC-AFFY cohort (partitioning around medoids (pam) clustering; “Pearson” distance; 1,000 iterations; from 2−7 clusters). The stability of the clusters was evaluated using the consensus matrix depicted as a dendrogram atop the heat map, the empirical cumulative distribution function (CDF) plot, and the delta area plot. To verify the repeatability and robustness of CCCRC, we used the “pamr.predict” function of the R package “pamr” (27) to classify the CRC samples based on the TME-related signature scores in the CRC-RNAseq cohort (seed = 11, threshold = 0.566). The TME-related signature scores were normalized by the Z-scores before performing “pamr.predict” analysis. PCOA based on euclidean distance was used to analyze the distribution of the CCCRC subtypes.

### Estimation of the TME cell abundance with other methods

The cell abundance of each sample was estimated based on the GEP using the microenvironment cell populations-counter (MCP-counter) algorithm (55) and the CIBERSORT (56) algorithm, both of which have been validated using the GEP of the corresponding cell populations and the degree of cellular infiltration estimated by immunohistochemistry. The MCP-counter algorithm estimated the cell abundance of 9 immune and stromal cell populations. The CIBERSORT algorithm, which applies the LM22 matrix, estimated the cell fraction of 22 immune cell populations. The ESTIMATE algorithm with default parameters was utilized to estimate the degree of infiltration of the total immune cells and stromal cells in the TME of each sample, as well as the tumor purity (57).

### Calculation of the other biological pathway enrichment scores

Human metabolism-related pathways were obtained from the Kyoto Encyclopedia of Genes and Genomes (KEGG) database (https://www.genome.jp/kegg/). The 1,660 genes assigned to 86 human metabolism-related pathways are listed in **Supplementary Table17**. 10 oncogenic signatures containing 331 genes and the terminally exhausted T cell signature were retrieved from a previously published study (58, 59) (**Supplementary Table17**). GSVA was performed to calculate the enrichment score of each signature for each sample of each cohort separately based on the relative GEP. To identify the potential differences in the biological functions of genes among CCCRC subtypes, gene set enrichment analysis (GSEA) was performed based on the gene signatures using R package “clusterprofiler” (60).

### Histopathological examination of the TCGA-CRC samples

A total of 616 TCGA CRC diagnostic hematoxylin and eosin (HE)-stained whole-slide images (WSIs) were downloaded from the TCGA data portal (March 2022) (https://portal.gdc.cancer.gov/), and the WSIs were examined blindly by two experienced pathologists. A total of 254 WSIs were included after removing the WSIs with poor quality and without views of the invasive margin (**Supplementary Table18**). According to the semi-quantitative pathological assessment of lymphocytes and their spatial location with malignant epithelial cells, the pathologist classified CRC into three immunophenotypes: “desert”, “excluded”, and “inflamed”, as previously described (32, 33). The inflamed phenotype was characterized by abundant lymphocytes in direct contact with malignant cells, the excluded phenotype was characterized by lymphocytes merely present in the stroma within or adjacent to the main tumor mass, and the desert phenotype was characterized by the lack of lymphocytes and stroma. We performed artificial intelligence (AI)-enabled spatial analysis of WSIs and developed a CRC-tissue classifier to identify eight tissue types: tumor, stroma, lymphocyte, normal colon mucosa, debris, adipose, mucin, and muscle, and quantified the abundances of the tumor, stroma, and lymphocytes in the core tumor (CT) region and the invasive margin (IM) region, respectively.

Our deep learning model (CRC-tissue classifier) consisted of two sequential parts: a muscle/non-muscle classifier that could distinguish each muscle patch in hematoxylin and eosin (H&E)-stained WSIs, and a seven-class tissue classifier that could classify seven tissue types: tumor, stroma, lymphocytes, normal colon mucosa, debris, adipose, and mucin. To develop the CRC-tissue classifier, we randomly selected 68,506 patches to train the muscle/non-muscle classifier and randomly selected 54,597 patches to train the seven-class tissue classifier, after combining the zenodo NCT-CRC-HE-100K dataset and the NCT-CRC-HE-100K dataset (https://zenodo.org/record/1214456#.YyRJGWB6RmM). Next, we evaluated the model on 4288 patches from 9 patients from the TCGA CRC datasets. The tissue regions were manually annotated by two experienced pathologists. The WSI tissue type prediction pipeline was as follows. First, the background was removed by the preprocessing steps. Second, the WSIs were segmented into non-overlapping image patches at a resolution of 0.5 μm/pixel (20 magnification). It is worth noting that if the WSI consisted of 40 magnifications, it was down-sampled to 20 magnifications. Next, the image patches were fed into the CRC-tissue classifier. If an image patch was determined to be non-muscle by the muscle/non-muscle classifier, it was fed into the multi-tissue classifier to predict its tissue class. We selected ResNet50 as the basic model architecture, adding one added full connection layer with ReLU as the activation function and 0.4 dropout: ReLU(x) = max (0, x), where x is the input of the ReLU function. Cross Entropy was selected as the loss function. During this experiment, we tested three model architectures, including ResNet50, vgg16, and Inception V3 for the multi-tissue classifier. According to the accuracy of seven tissues (tumor, stroma, lymphocytes, normal colon mucosa, debris, adipose, and mucin) in the CRC-7k dataset, the performance of ResNet50 was the best, which was the reason we selected ResNet50 as the basic model architecture.

After recognizing the CRC tissue types by our deep learning model automatically, we quantified the abundances of the tumor, lymphocytes, and stroma in the core tumor (CT) region and the invasive margin (IM) region. The quantification pipeline consisted of four steps. First, we used the open source software QuPath-0.3.2 (https://qupath.github.io/) to delineate the CT and IM region. The IM region was defined as 500mm outside the CT region (61). The CT and IM regions were manually annotated by two experienced pathologists to reduce bias. Second, the abundances of the tumor, lymphocytes, and stroma in each WSI were quantified with an area ratio of their area. Finally, we calculated the mean abundances of the tumor, lymphocytes, and stroma in each WSI. A total of 254 TCGA-CRC WSIs were quantified.

### Acquisition of signatures associated with the immune checkpoint blockade (ICB) therapy response

The Tumor Immune Dysfunction and Exclusion (TIDE) score was calculated using GEP, and it was used to evaluate the degree of T cell dysfunction and T cell exclusion (62). The higher the score, the later the dysfunction stage of T cells or the higher the degree of T cell exclusion. The gene expression average of all samples in each cohort was used as the normalized control and the normalized gene expression matrix was uploaded to the TIDE website (http://tide.dfci.harvard.edu/).

### Acquisition and processing of CRC multi-omics data

Masked somatic mutation data (n = 571 samples), masked copy number segment data (n = 609 samples) and DNA methylation beta-values (Illumina human methylation 450) (45 normal samples and 390 tumor samples) were download from the TCGA data portal (March 2022) (https://portal.gdc.cancer.gov/). The liquid chromatography-tandem mass spectrometry (LC-MS/MS)-based proteomic data for the TCGA CRC samples (n = 88 samples) were obtained from a previously published study (63). The R package “maftools 2.6.05” with default parameters was used to analyze the somatic mutation data. Synonymous mutations were regarded as wild-type, and genes with mutation rates <5% were excluded. Nonsynonymous mutations were used to calculate tumor mutation burden (TMB). Somatic copy number alterations (SCNA) defined by the GISTIC2.0 module on the GenePattern website (https://www.genepattern.org/), including arm-level gain (1), and high amplification (2), diploid/normal (0), arm-level deletion (−1), and deep deletion (−2). The CINmetrics algorithm was used to calculate chromosomal instability signature (CIN), including SCNA count and fraction of the genome altered (FGA), which was proposed by Vishaloza et al. (https://rdrr.io/github/lasseignelab/CINmetrics/) based on previously published studies (64–66). If somatic mutation events or SCNAs occurred in one or more genes in the oncogenic pathway, the tumor sample was considered altered in a given pathway. The microsatellite (MSI) status was obtained from the CMS website (https://www.synapse.org/#!Synapse:syn2623706). Tumor neoantigen signature were obtained from a previously published study (67). The prevalence of somatic mutation events or SCNAs was compared among CCCRC cases using Fishers exact test or chi-square test. For the DNA methylation data, probes located in promoter CpG islands were extracted, including TSS200, 1stExon, TSS1500, and 5′UTR. The probes detected on X and Y chromosomes or any probe with NA value were removed. For genes with multiple probes mapped to the promoter, the median beta-value was calculated as the degree of gene methylation. The beta-value difference was defined as the difference between the mean beta value of each CCCRC sample and normal samples, and Wilcoxon rank-sum test was used to test whether the difference was statistically significant. *P*-values were adjusted for multiple comparisons by the FDR method. Differentially methylated genes (DMGs) between normal and CRC samples were defined as |mean beta value| <0.2 in normal samples, |mean beta value| >0.5 in CRC samples, and FDR <0.05. DMGs between CCCRC subtypes were defined as FDR <0.001. To identify differentially expressed genes (DEGs) between CCCRC subtypes in the CRC-AFFY and CRC-RNAseq cohorts, the “limma” package was used with FDR <0.001. Wilcoxon rank-sum test was used to identify differentially expressed proteins (DEPs) with P-values <0.05 between CCCRC subtypes.

### Regulon analysis

The R package “RTN” was used to reconstruct the transcriptional regulatory networks of regulons (68), including 31 transcription factors and 82 chromatin remodeling genes, that were associated with CRC (69, 70) (**Supplementary Table19**). Mutual information and Spearman’s correlation analysis were utilized to infer the possible associations between a regulator and all possible targets from the GEP, and the permutation algorithm was used to eliminate associations with an FDR >1×10^-5^. Unstable associations were removed by bootstrap analysis (n = 1,000), and the weakest association in triangles consisting of two regulators and common targets were eliminated by the data processing inequality algorithm. Two-tailed gene set enrichment analysis was used to calculate the regulon activity score for each sample.

### Publicly available CRC classification systems

To classify CRC samples into different CRC subtypes according to the previously published gene classifier, gene lists for the five classifiers were extracted from relevant publications and summarized (**Supplementary Table20**), including Budinska subtypes (6), De Sousa subtypes (7), Roepman subtypes (9), and Sadanandam subtypes (10).The nearest template prediction (NTP) algorithm was employed to classify the samples and to generate an FDR to assess the classification robustness. For NTP implementation, we screened genes that were specifically and positively associated with one subtype according to the screening strategies of a previously published study (71).

### Bulk RNAseq and scRNAseq data processing of the GSE108989 dataset

A total of 12 CRC samples with bulk RNAseq and scRNAseq data were obtained from the GSE108989 dataset (https://www.ncbi.nlm.nih.gov/geo/query/acc.cgi?acc=GSE108989) (72). To identify the CCCRC subtypes, bulk RNAseq with transcripts per million (TPM) was further log2-transformed, and GSVA was performed to calculate the signature score of each TME-related signature in each sample based on the GEP. The “pamr.predict” algorithm was used to classify CRC samples into four CCCRC subtypes based on the TME-related signatures (seed = 11, threshold = 0.566). For scRNAseq data processing, the raw gene expression data were normalized and selected according to the following criteria: cells with >200 genes and <7,000 genes and <20% of mitochondrial gene expression in UMI counts, which was determined using the Seurat R package. Counts of the filtered matrix for each gene were normalized to the total library size with the Seurat “NormalizeData” function. The “FindVariableGenes” function was used to identify 2,000 hypervariable genes for unsupervised clustering. Next, each integrated feature was centered to a mean of zero and scaled by the standard deviation with the Seurat “ScaleData” function. The “RunPCA” function was used for PCA. We identified diverse T cell clusters using the “FindClusters” function, and set the resolution parameter to 0.5. Each cell cluster was compared to the other clusters by the “FindAllMarkers” function to identify DEGs (only pos: TRUE, min.PCt: 0.25, logFc.threshold: 0.25). Cell annotation was carried out by consulting the latest cell marker databases, such as CellMarker (https://www.biolegend.com/en-us/cell-markers) and PanglaoDB (https://ngdc.cncb.ac.cn/databasecommons/database/id/6917), combined with a previously published study (72). To define the feature genes for each CCCRC subtype, differential expression analysis between CCCRC subtypes was performed using the “FindMarkers” function. FDR <0.05 were considered statistically significant.

### Collection and processing of therapy-associated datasets

Therapy-associated datasets were used to explore the treatment strategies for each CCCRC subtype. Gene expression profiles of GSE103479 and GSE104645 datasets were downloaded from the GEO database. If the gene symbol was annotated with multiple probes, the median value was used as the expression of the gene. The clinical data of the GSE104645 dataset was obtained from the supplementary table of a study by Okita et al. (73). The GSE103479 dataset contained 156 stage II and III CRC patients with or without 5-fluorouracil (5FU)-based chemotherapy. The GSE104645 dataset contained 193 mCRC patients treated with chemotherapy, a combination of chemotherapy and bevacizumab, or anti-EGFR therapies. The available RNAseq expression dataset of patients treated with anti-PD-1 therapy was also downloaded. The Gide (PRJEB23709) dataset was downloaded, and the raw fastq files was re-analyzed. The RNA reads were aligned using STAR v2.5.3 and quantified as TPM using RSEM v1.3.0 and log_2_-transformed. Ensembl IDs were annotated into gene symbols using GENCODE v36. The gene expression profiles of Hugo (GSE78220) and Jung (GSE135222) datasets and the corresponding clinical data were downloaded from the GEO database, and the FPKM values were converted to log2-transformed TPM values. We obtained the gene expression data (n = 348) of urothelial carcinoma patients in the IMvigor210 dataset treated with anti-PD-L1 therapy and the corresponding clinical data using R package “IMvigor210CoreBiologies 1.0.0” (IMvigor210 dataset), and the count values were converted to log_2_-transformed TPM values. To reduce batch effects and tissue-type-specific effects, we first performed GSVA analysis of the TME-related signatures in each dataset, and the signature scores were normalized by Z-scores before using the “pamr” algorithm. Next, we used the “pamr” algorithm to classify the samples into the four CCCRC subtypes based on the TME-related signatures in each dataset (seed = 11, threshold = 0.566). Detailed information on the sample size and the corresponding treatment data of the therapy-associated datasets are summarized in **Supplementary Table21**.

To explore the treatment for each CCCRC subtype using cancer cell line drug-sensitivity experiments, we developed a pre-clinical model based on subtype-specific, cancer cell-intrinsic gene markers according to a previously published study (41). The CCCRC subtype-specific mRNA genes (log2 (fold change) >0 and FDR <0.05) was determined by R package “limma” based on RMA normalization data in the CRC-AFFY cohort. The gene expression of human CRC tissues versus patient-derived xenografts in the GSE35144 dataset by the R package “limma” was used to remove those genes associated with stromal and immune components. DEGs with FDR >0.5 and log2 (fold change) <2 were considered as cancer cell-intrinsic genes. A total of 71 human CRC cell lines with RNAseq data (log2TPM) was obtained from the Genomics of Drug Sensitivity in Cancer (GDSC) database (https://depmap.org/portal/download/all/), 43 of which had drug sensitivity results. RNAseq data for 71 human CRC cell lines was used to further determine the cancer cell-intrinsic genes and genes among the top 25% within (i) the 10−90 % percentile range of the largest expression values and (ii) the highest expression in at least three samples. The subtype-specific genes and cancer cell-intrinsic genes were intersected and further screened by the Boruta importance test to generate the gene list for developing the pre-clinical model. The GSE13067, GSE13294, GSE33113, GSE37892, GSE38832, and GSE39582 datasets were combined as the training set and the GSE14333 and GSE17536 datasets were used as the validation set, separately. The GEP of CRC cell lines was normalized by the “quantileNormalizeByFeature” function in the package of “FSQN” (74). The random forest algorithm (RF), support vector machine algorithm (SVM), eXtreme Gradient Boosting (xgboost) algorithm, and logistic regression algorithm was used to develop the pre-clinical models. The accuracy, F1 values, and AUC values were computed to evaluate the performance of the pre-clinical models. We used the pre-clinical model with best predictive performance to classify 71 human CRC cell lines into four CCCRC subtypes and compared the differences of the area under the receiver operator characteristics curve (AUC) drug responses among the CCCRC subtypes.

### Discovery and validation of the single-sample gene classifier

Considering that the current transcriptomic data were mostly based on next-generation sequencing platforms, we constructed and validated a single-sample model to identify CCCRC subtypes based on CRC-RNAseq cohort. The R package “limma” was used to determine subtype-specific mRNA genes (log2 (fold change) >0 and FDR <0.05) based on the “voom” transformation with quantile normalization data in the CRC-RNAseq cohort. The Boruta importance test was further performed to screen subtype-specific mRNA genes. The CRC-RNAseq cohort was randomly divided into the training set and the validation set at a ratio of 3:7. The gene expression data was normalized by the Z-scores before model training and could be applied to a single-sample setting. The single-sample gene classifiers were trained with the random forest algorithm (RF), support vector machine algorithm (SVM), eXtreme Gradient Boosting (xgboost) algorithm, and logistic regression algorithm using the subtype-specific genes. We also validated the gene classifier in TCGA and CPTAC dataset. The accuracy, F1 values, and AUC values were computed to evaluate the predictive performance of the single-sample gene classifiers.

### Statistical analyses

All statistical analyses were conducted by R 4.0.2 software. Statistical significance of the comparisons for continuous variables and categorical variables was assessed by the Wilcoxon rank-sum test or Kruskal-Wallis test and Fisher’s exact test or chi-square test, respectively. Correlations between variables were estimated by Spearman’s correlation analysis or Pearson’s correlation analysis. Patients were divided into either high or low gene expression groups by the best cutoff calculated by the R package “survminer”. The Kaplan-Meier method with log-rank test was utilized to generate the survival curves. Univariate and multivariate Cox proportional hazard regression analyses were performed to generate 95% confidence intervals (CIs) and hazard ratios (HRs). Two-sided *P*-values <0.05 were considered statistically significant.

## Acknowledgments

This work was supported in part by research grants from the National Key R&D Program of China (Grant No. 2017YFC1309002) to L.L. We thank TCGA and GEO databases and the cBioportal website for free use. We thank Nanjing Simcere Medical Laboratory Science Co., Ltd. and Jiangsu Simcere Diagnostics Co., Ltd., and all the members of its AI and bioinformatics team, for generously sharing their experience and codes.

## Supplementary Figure Legends

**Figure S1.**
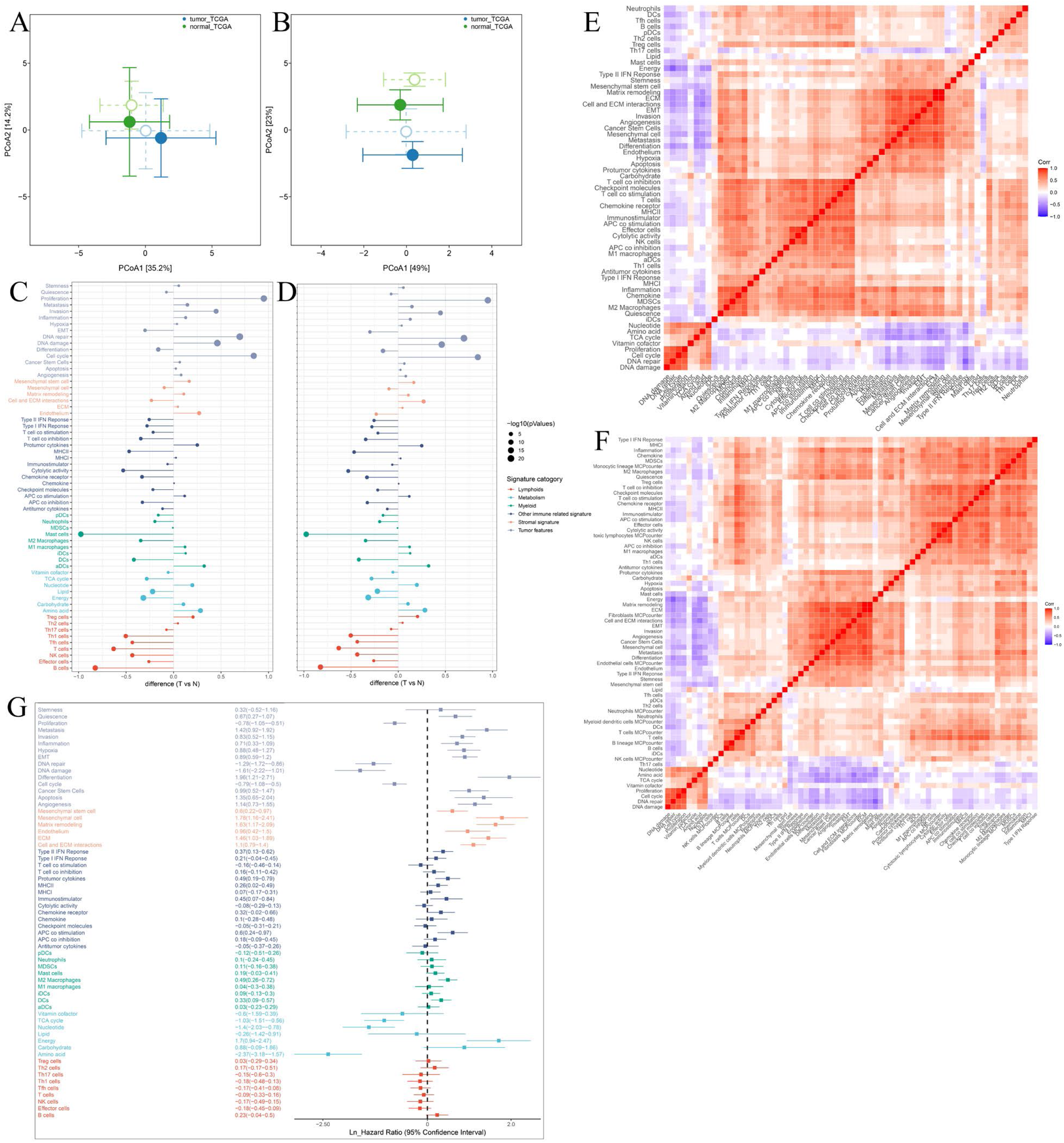
Establishment of TME gene expression panel. **A, B.** PCOA shows that the CRC samples could be distinguished from the normal samples by the TME-related signatures (**A**) and the signatures of the functional states of tumor cells and cancer stem cells (**B**) in the GSE39582 and TCGA cohorts. **C, D.** Difference analysis of TME signature scores between tumor (T) and normal (N) tissues in the GSE39582 cohort (**C**) and the TCGA cohort (**D**). **E.** Pearson’s correlation analysis of the TME-related signatures show four major patterns bound by positive correlations in the CRC-AFFY cohort. **F.** Heat map of Pearson’s correlation analysis of the 61 TME-related signatures and the other TME-related signatures quantified by the MCP-counter algorithm in the CRC-AFFY cohort. **G.** Univariate cox analysis shows the ability of each TME signature to predict progression-free survival in the CRC-AFFY cohort.

**Figure S2.**
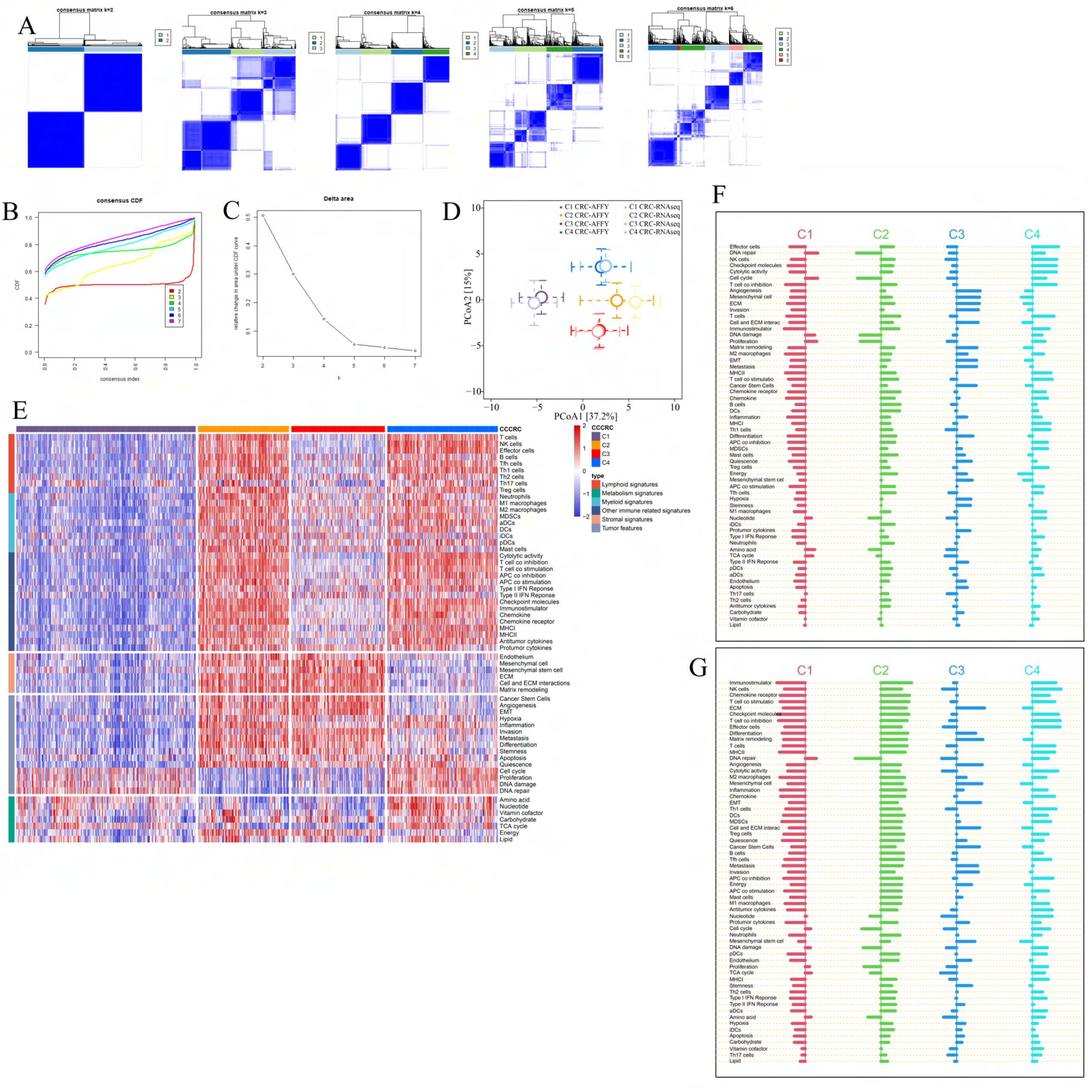
Comprehensive characterization of colorectal cancer (CCCRC). **A.** Consensus matrices heat map (k = 2 to 6). **B.** Empirical cumulative distribution function (CDF) plot. **C.** Delta area plot. **D.** Principal coordinate analysis of Euclidean distances calculated using the scores of 61 TME-related signatures in the CRC-AFFY (dark colors) and CRC-RNAseq (light colors) cohorts. Circles and error bars represent the mean and the standard errors of the mean, respectively. **E.** Heat map of 725 CRC patients in the CRC-RNAseq cohort classified into four distinct TME subtypes based on the 61 TME-related signatures. **F, G.** Shrunken differences d′ik for the 61 TME-related signatures having at least one nonzero difference in the CRC-AFFY cohort (**F**) and the CRC-RNAseq cohort (**G**).

**Figure S3.**
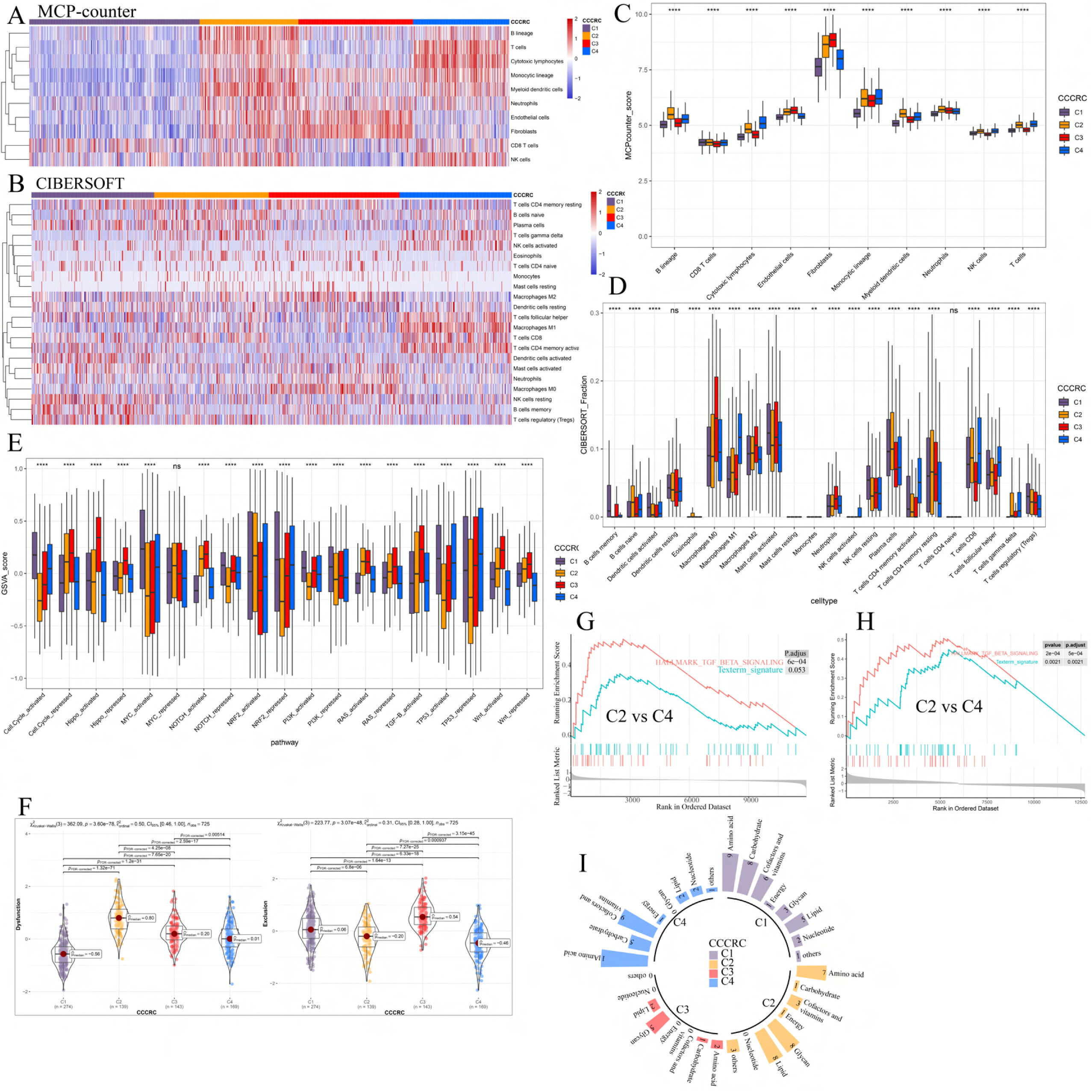
Differences in the TME components obtained from MCP-counter, CIBERSORT, and the ESTIMATE algorithm among the CCCRC subtypes in the CRC-AFFY cohort. **A, B.** Heat map of the TME-related signature scores derived from the MCP-counter (**A**) and CIBERSORT (**B**) algorithm. **C-E.** Box plots show differences in the TME-related signature scores derived from the MCP-counter (**C**), CIBERSORT (**D**), and the GSVA (**E**) algorithm among the CCCRC subtypes. **F.** Differences in T cell dysfunction and T cell exclusion scores between four CCCRC subtypes were analyzed based on the gene expression profiles in CRC-RNAseq cohort. **G, H.** Gene set enrichment analysis (GSEA) of the terminally exhausted CD8+ T cell signature (Texterm signature) and the TGFB signaling signature between C2 and C4 subtypes in the CRC-AFFY cohort (**G**) and CRC-RNAseq cohort (**H**). **I.** Circle bars display significant differences in metabolic reprogramming among the four CCCRC subtypes. *p value < 0.05; **p value < 0.01; ***p value < 0.001; ****p value < 0.0001.

**Figure S4.**
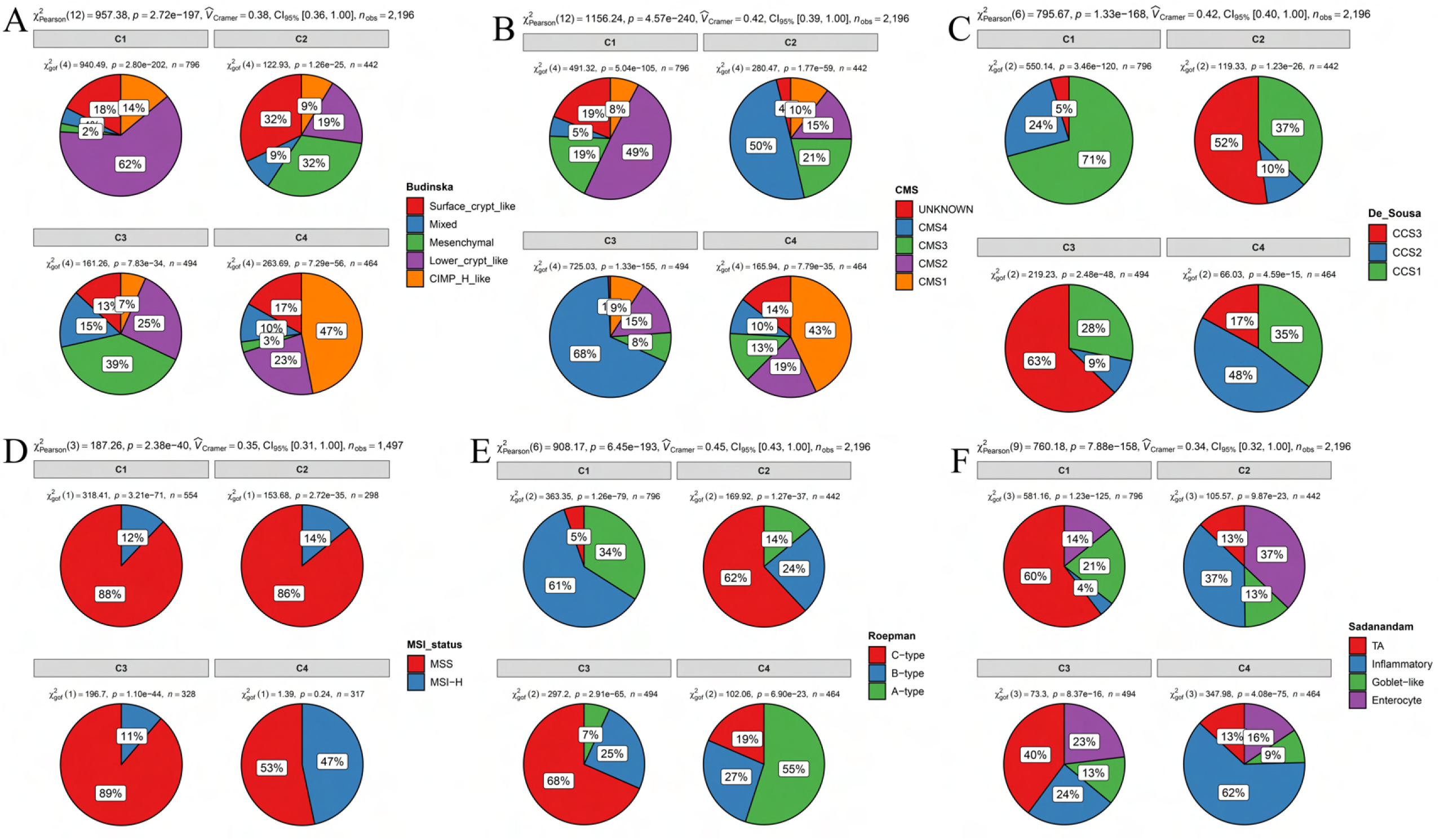
Overlap of the CCCRC subtypes with published CRC molecular subtypes in the CRC-AFFY and CRC-RNAseq cohorts, including Budinska subtypes. **A,** Consensus molecular subtypes (CMS). **B,** De Sousa subtypes. **C,** Microsatellite instability (MSI) status (high microsatellite instability [MSI-H]). **D,** Microsatellite stability (MSS). **E,** Roepman subtypes. **F,** Sadanandam subtypes.

**Figure S5.**
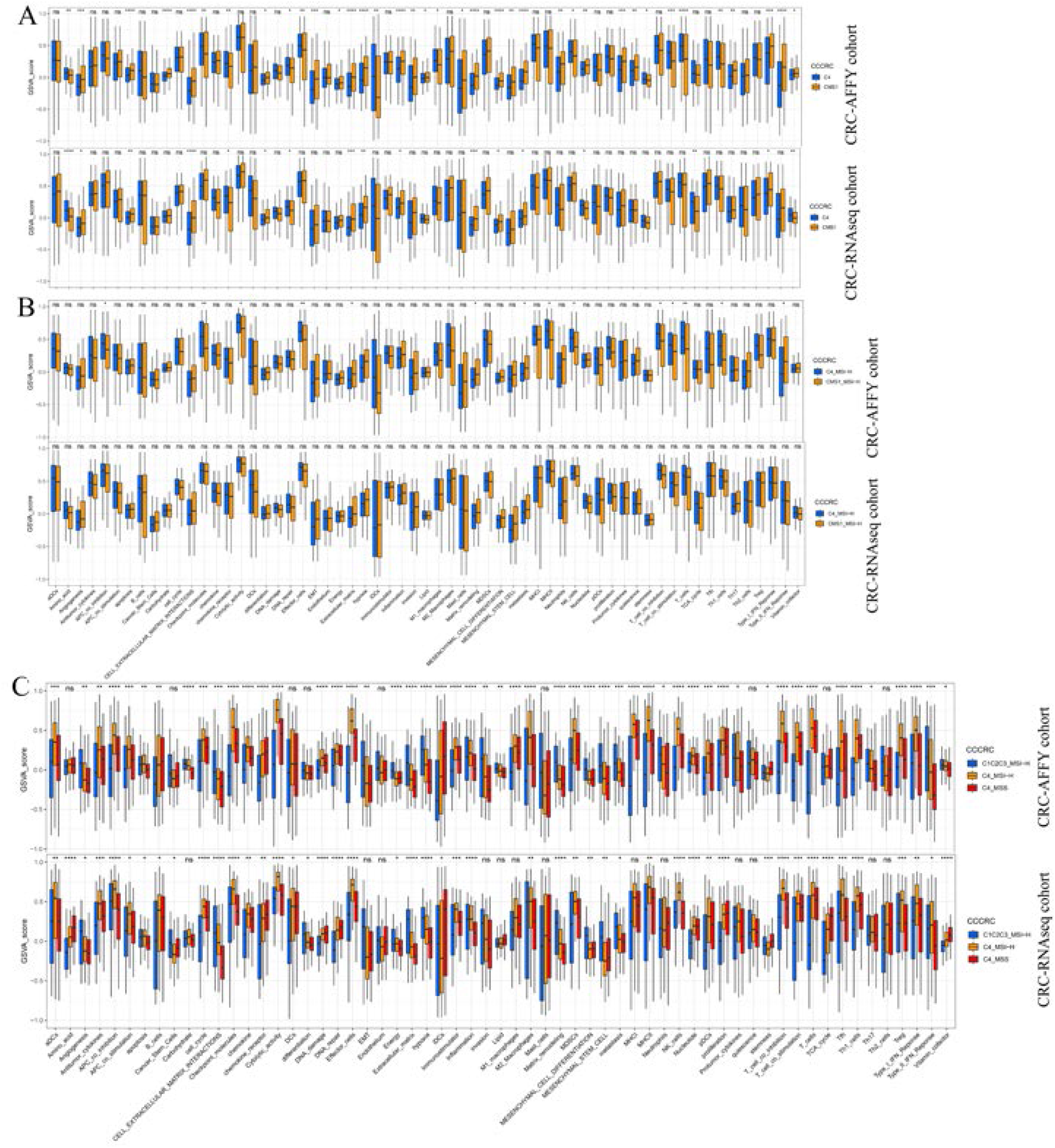
Box plots show differences in the TME-related signature scores between the C4 and CMS1 subtypes. **A,** Between the C4 subtype with MSI-H and the CMS1 subtype with MSI-H. **B,** Between the C4 subtype with MSI-H, the C4 subtype with MSS, and the other CCCRC subtypes with MSI-H. **C,** MSI-H, high microsatellite instability. *p value < 0.05; **p value < 0.01; ***p value < 0.001; ****p value < 0.0001.

**Figure S6.**
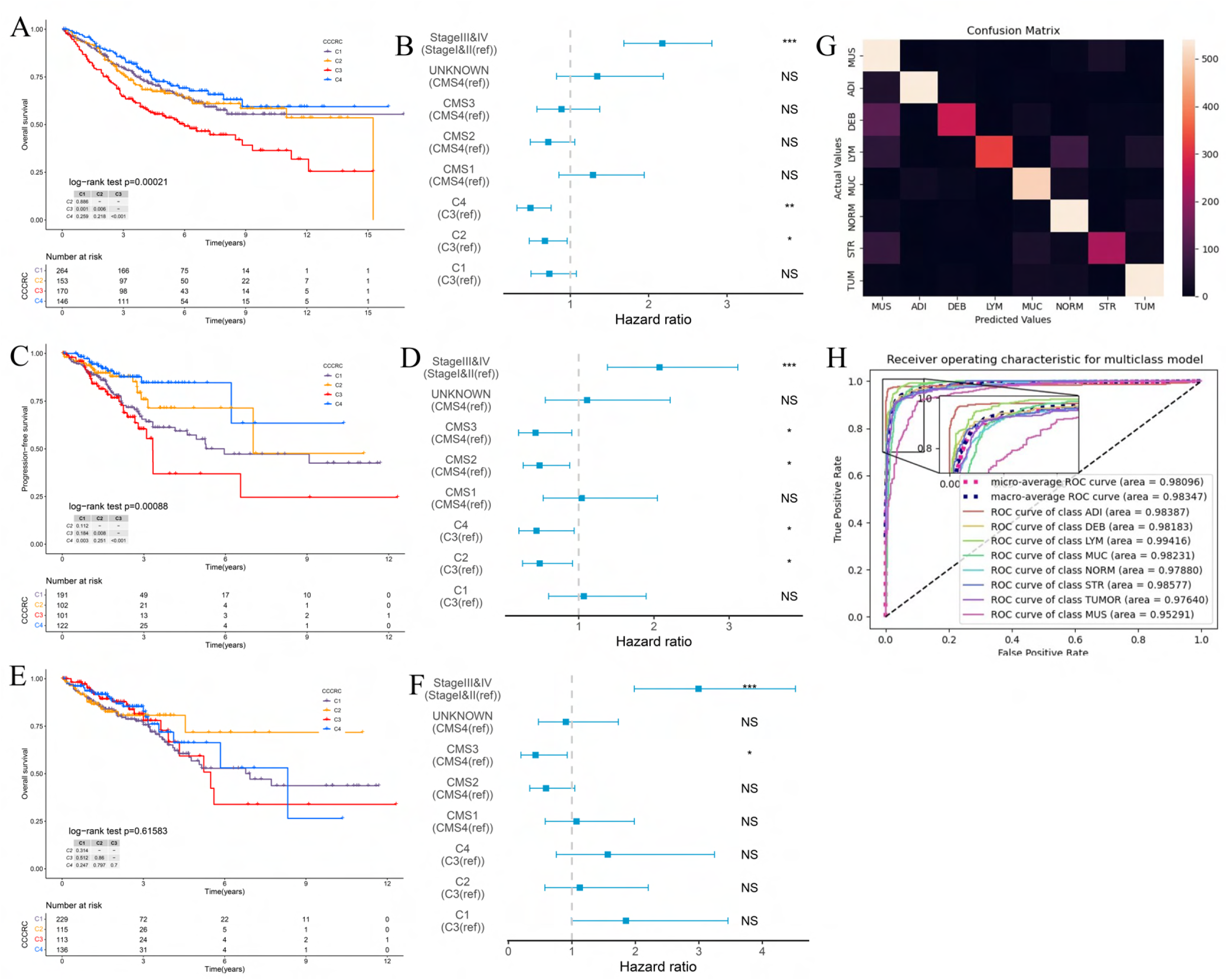
Survival analyses of the CCCRC subtypes. **A,** Kaplan-Meier method with log-rank test of overall survival (OS) among the four CCCRC subtypes in the CRC-AFFY cohort. **B,** Forest plot of multivariate Cox proportional hazard regression analysis for OS after adjusting for TNM stage and CMS subtype in the CRC-AFFY cohort. **C, D,** Kaplan–Meier method (**C**) and multivariate Cox proportional hazard regression analysis (**D**) of progression-free survival (PFS) among the four CCCRC subtypes in the CRC-RNAseq cohort. **E, F**, Kaplan-Meier method with log-rank test (**E**) and multivariate Cox proportional hazard regression analysis (**F**) of OS among the four CCCRC subtypes in the CRC-RNAseq cohort. The hazard ratios are shown with 95% confidence intervals. *p value < 0.05; **p value < 0.01; ***p value < 0.001; NS, p value > 0.05. **G.** Confusion matrix shows overlapping numbers of predicted tissues and actual tissues. **H.** AUC curves show performance of the CRC-multiclass model on the TCGA-CRC dataset.

**Figure S7.**
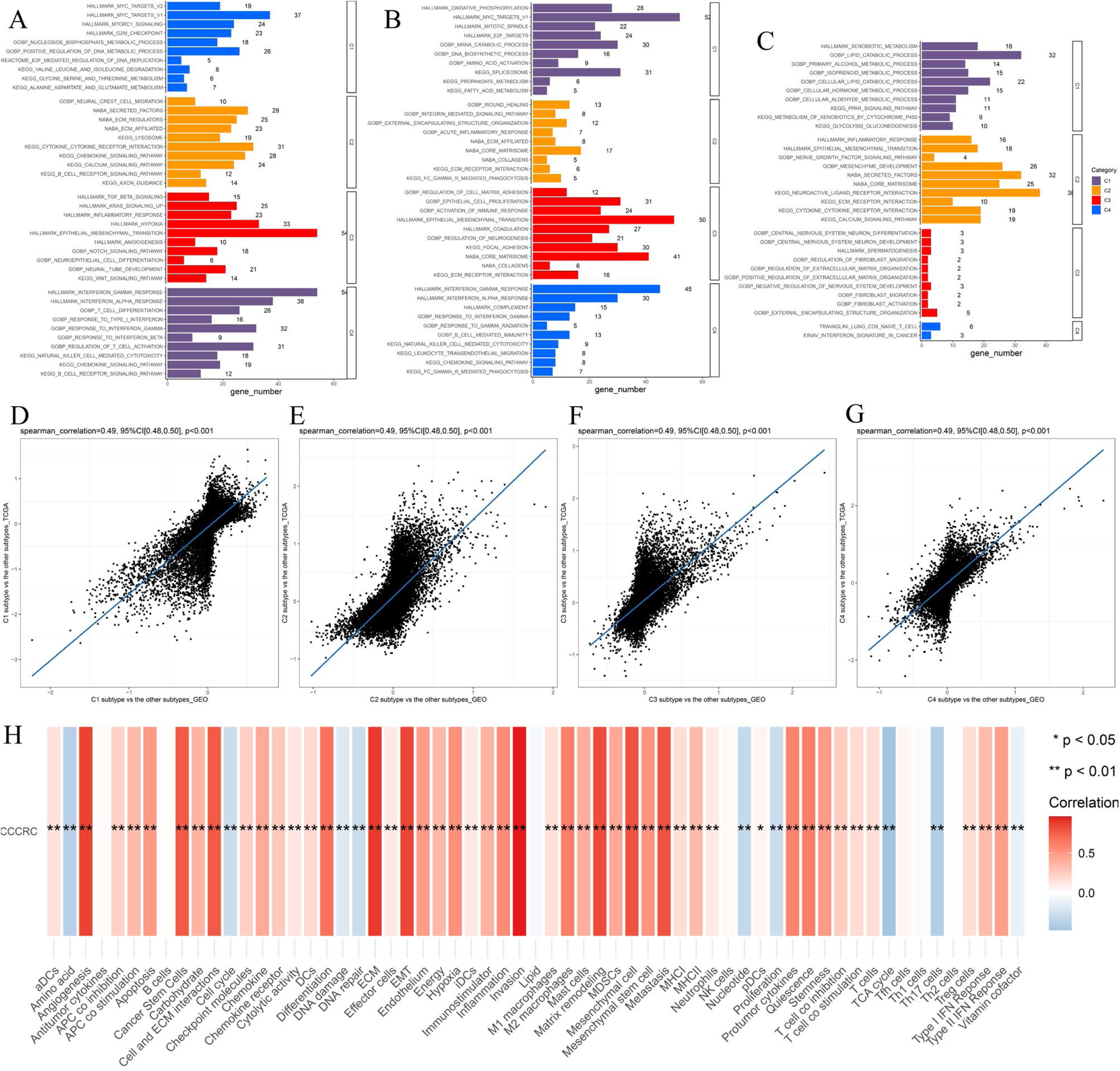
Biological characterization of the CCCRC subtypes based on multi-omics data. **A-C,** Significantly enriched gene sets among the CCCRC subtype-specific upregulated genes, CCCRC subtype-specific downregulated methylation genes, and CCCRC subtype-specific upregulated proteins. **D-G,** Scatter plots show gene expression log2-fold changes for all genes among the four CCCRC subtypes (C1 subtype vs the other subtypes, **D**; C2 subtype vs the other subtypes, **E**; C3 subtype vs the other subtypes, **F**; and C4 subtype vs the other subtypes, **G**) in the CRC-AFFY cohort and the CRC-RNAseq cohort. **H,** Relationship between CCCRC scores and TME-related signature scores.

**Figure S8.**
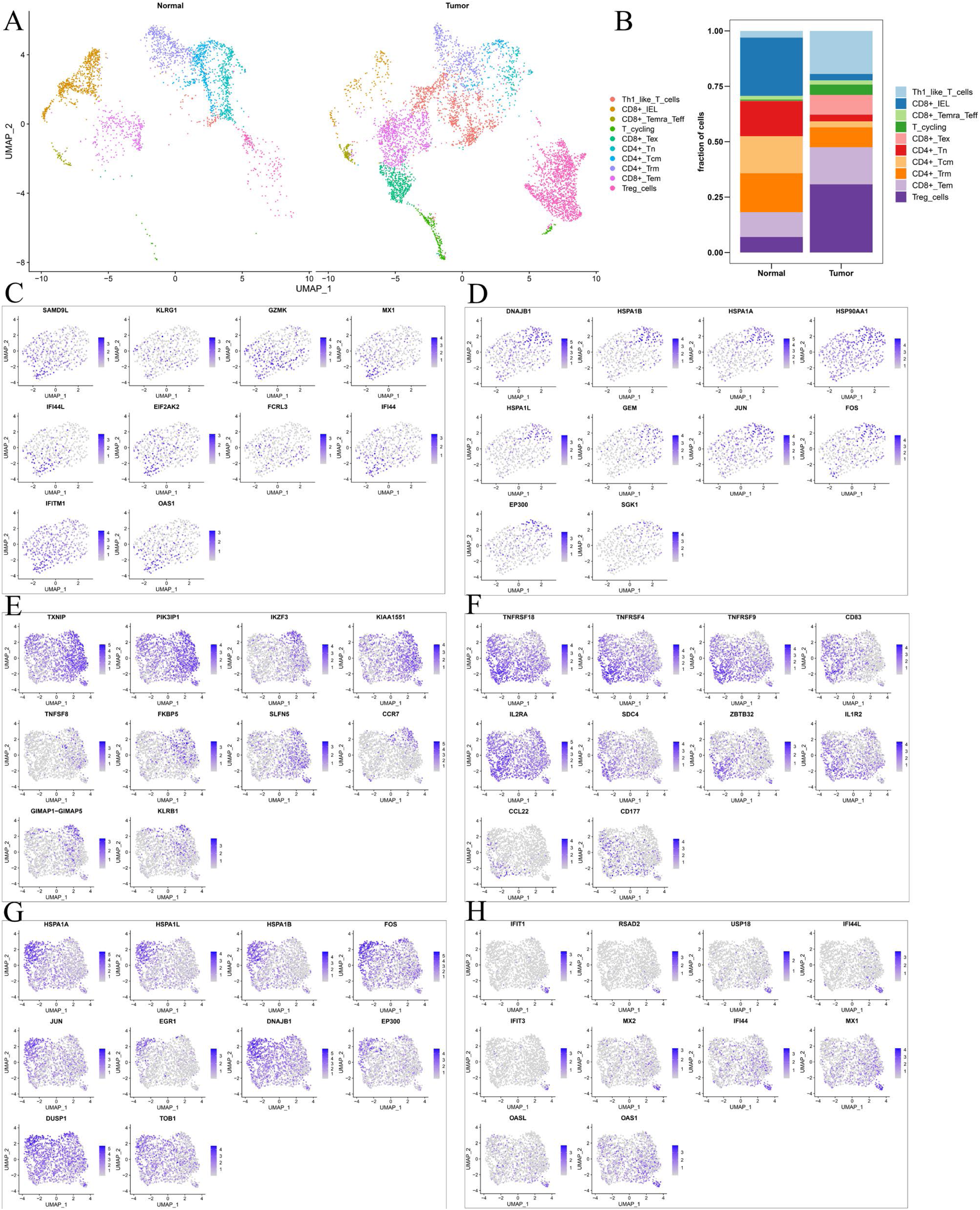
**A,** UMAP shows the composition of T cells colored by cluster in tumor and normal tissues. **B,** UMAP shows the composition of T cells colored by cluster and classified by CCCRC subtype in tumor and normal tissues. **C-H,** The tSNE visualized plot shows the expression of the top 10 marker genes for KLRG1+ CD8+ Tex (**C**), HSPA1B+ CD8+ Tex cells (**D**), TXNIP+ Treg cells (**E**), TNFRSF4+ Treg cells (**F**), HSPA1A+ Treg cells (**G**), and IFIT1+ Treg cells (**H**).

**Figure S9.**
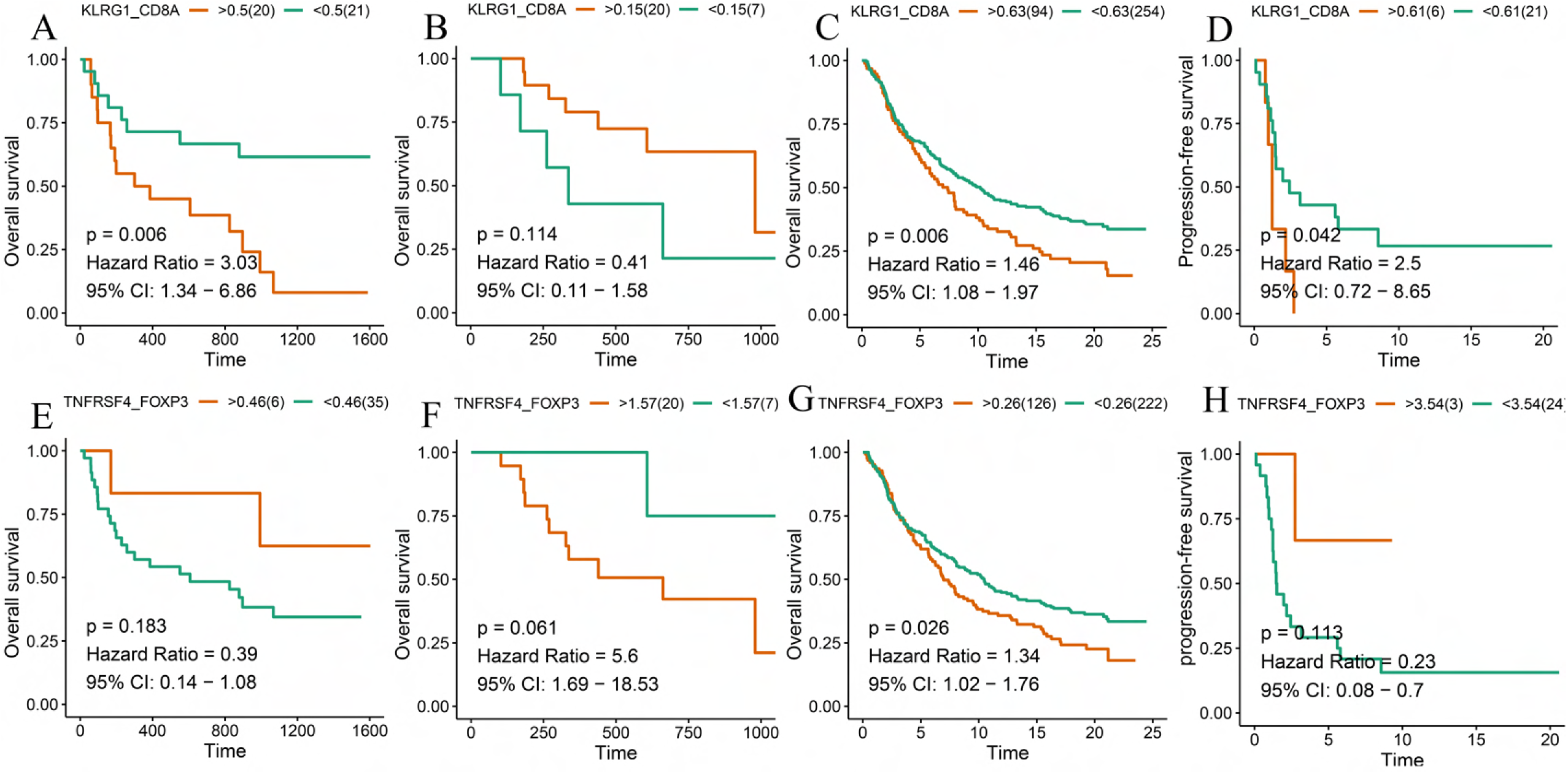
**A-D,** Kaplan-Meier method with log-rank test of overall survival (OS) and progression-free survival (PFS) between the high ratio of KLRG1-to-CD8A expression and the low ratio of KLRG1-to-CD8A expression in Gide (**A**), Hugo (**B**), Jung (**C**), and IMvigor210 (**D**) datasets. **E-H,** Kaplan-Meier method with log-rank test of OS and PFS between the high ratio of TNFRSF4-to-FOXP3 expression and the low ratio of TNFRSF4-to-FOXP3 expression in Gide (**E**), Hugo (F), Jung (**G**), and IMvigor210 (**H**) datasets.

**Figure S10.**
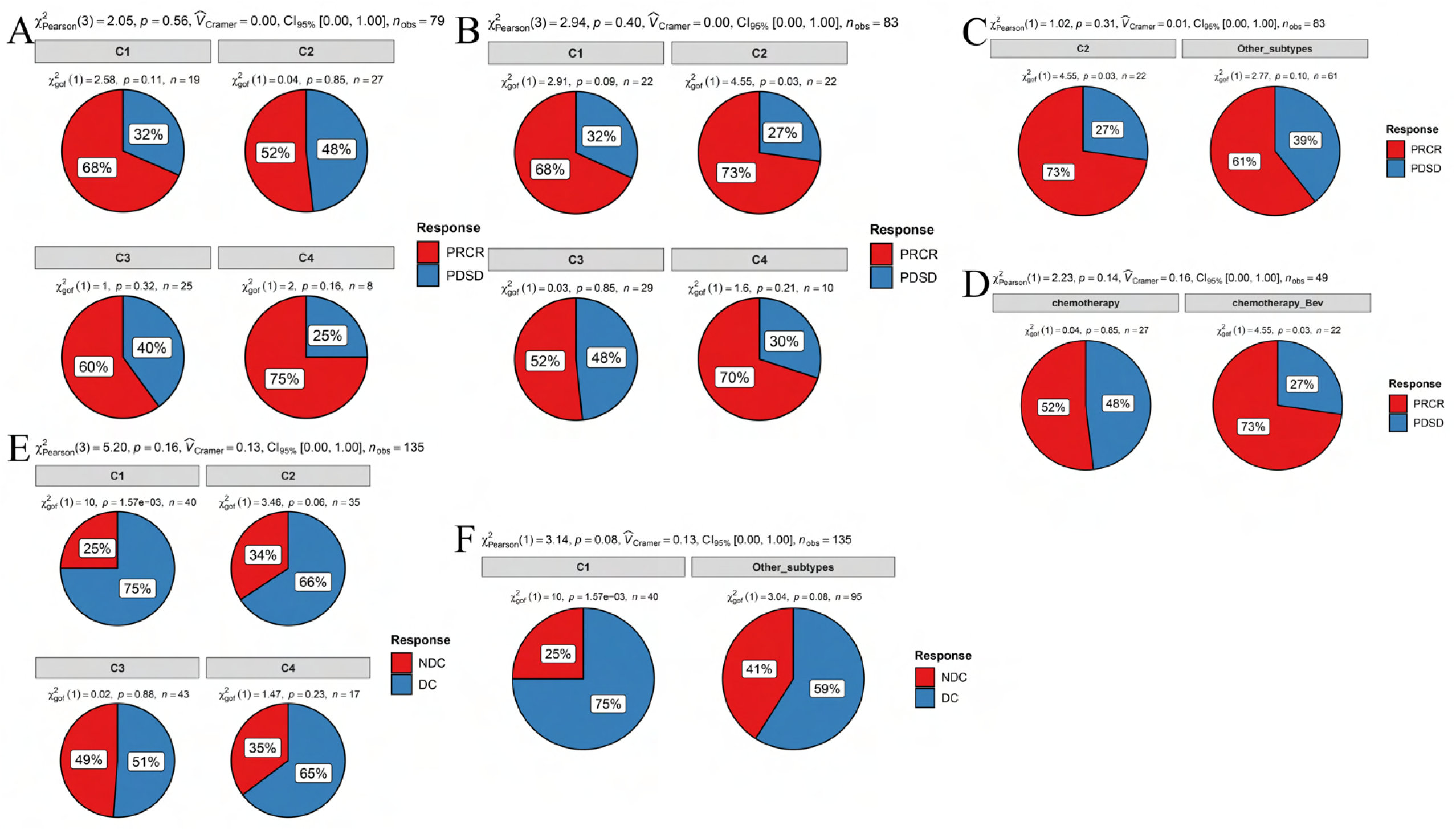
**A,** Pie chart shows the differences in the proportions of responses to chemotherapy among the four CCCRC subtypes in the GSE104645 dataset. **B, C,** Pie chart shows the differences in the proportions of responses to a combination of chemotherapy and bevacizumab among the four CCCRC subtypes (**B**) and between the C2 subtype and the other subtypes (**C**) in the GSE104645 dataset. **D,** Pie chart shows the differences in the proportions of responses to chemotherapy plus bevacizumab versus responses to chemotherapy in the C2 subtype of the GSE104645 dataset. **E,** Pie chart shows the differences in the proportions of the disease control rate (DCR) of anti-EGFR therapy among the four CCCRC subtypes in the GSE104645 dataset. **F,** Pie chart shows the differences in the proportions of responses to anti-EGFR therapy between the C2 subtype and the other subtypes in the GSE104645 dataset. PRCR, partial response and complete response; PDSD, progressive disease and stable disease; DC, disease control; NDC, no disease control.

**Figure S11.**
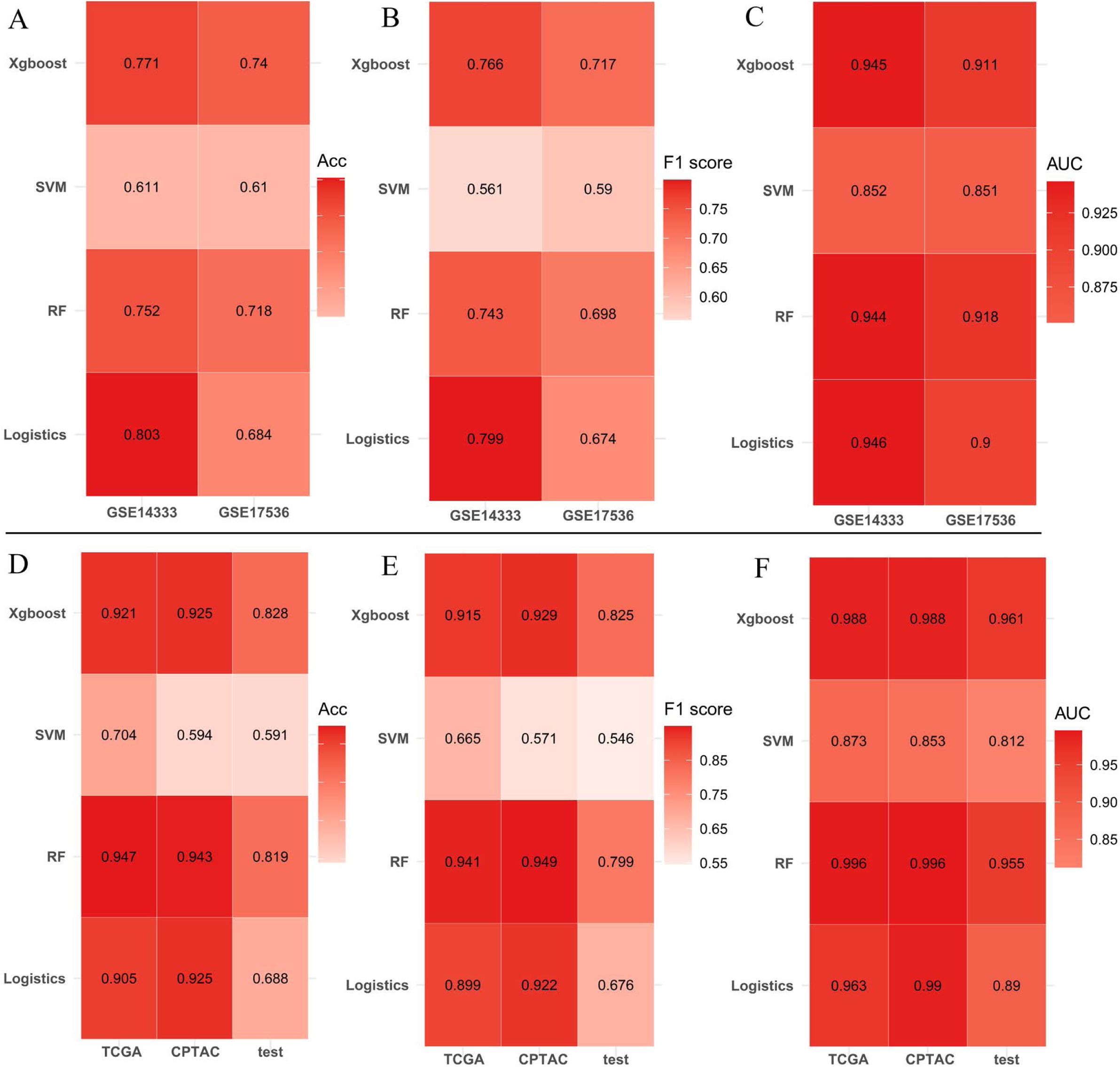
Establishment of machine learning model. **A-C.** The pre-clinical model was constructed using the random forest algorithm (RF), support vector machine algorithm (SVM), extreme gradient boosting (xgboost) algorithm, logistic regression algorithm. Accuracy (**A**), F1 score (**B**), and AUC value (**C**) were computed to evaluate the performance of the models. **D-F.** The single-sample gene classifier was constructed using the RF, SVM, xgboost algorithm, logistic regression algorithm. Accuracy (**D**), F1 score (**E**), and AUC value (**F**) were computed to evaluate the performance of the classifiers.

